# Screening of the Key Genes and Signaling Pathways for Schizophrenia Using Bioinformatics and Next Generation Sequencing Data Analysis

**DOI:** 10.1101/2023.10.24.563759

**Authors:** Basavaraj Vastrad, Chanabasayya Vastrad

**Author notes:** Chanabasayya Vastrad, Chanabasava Nilaya, Bharthinagar, Dharwad 580001, Karanataka, India.

## Abstract

Schizophrenia is thought to be the most prevalent chronic psychiatric disorder. Numerous proteins have been identified that are associated with the occurrence and development of schizophrenia. This study aimed to identify potential core genes and pathways involved in schizophrenia, through exhaustive bioinformatic and next generation sequencing (NGS) data analyses using GSE20966 NGS data of neural progenitor cells and neurons obtained from healthy controls and patients with schizophrenia. The NGS data were downloaded from the Gene Expression Omnibus database. NGS data was processed by the DESeq2 package in R software and the differentially expressed genes (DEGs) were identified. Gene Ontology (GO) enrichment analysis and REACTOME pathway enrichment analysis were carried out to identify potential biological functions and pathways of the DEGs. Protein-protein interaction (PPI) network, module, miRNA-hub gene regulatory network and TF-hub gene regulatory network analysis were performed to identify the hub genes, miRNA and TFs. Potential hub genes were analyzed using receiver operating characteristic (ROC) curves in the R package (pROC). In this investigation, an overall 955 DEGs were identified: 478 genes were remarkably up regulated and 477 genes were distinctly down regulated. These genes were enriched for GO terms and pathways mainly involved in the multicellular organismal process, GPCR ligand binding, regulation of cellular process and amine ligand-binding receptors. MYC, FN1, CDKN2A, EEF1G, CAV1, ONECUT1, SYK, MAPK13, TFAP2A and BTK were considered the potential hub genes. miRNA-hub gene regulatory network and TF-hub gene regulatory network were constructed successfully. On the whole, the findings of this investigation enhance our understanding of the potential molecular mechanisms of schizophrenia and provide potential targets for further investigation.

## Introduction

Schizophrenia is a chronic brain disease in which imbalance in serial neurotransmitters, such as dopamine and glutamate [1]. Schizophrenia affects around 1% of the world population that causes a severe health burden [2]. The main features of progression of schizophrenia are delusions, hallucinations, thought disorders, anhedonia, avolition, social withdrawal, poverty of thought and cognitive dysfunction [3]. Schizophrenia, one of the major components of the heterogeneous psychiatric disorder, is closely associated with a variety of diseases such as cardiovascular diseases [4], neurodegenerative diseases [5], infections [6], obesity [7], diabetes mellitus [8] and hypertension [9]. Although there are extensive studies on the molecular mechanism in schizophrenia progression, the causes of schizophrenia is still not clear. The occurrence and progression of schizophrenia are correlated with multiple factors from the point of view of science and research, for instance, genetic and environmental factors [10]. The causes and the underlying molecular mechanisms, discovering molecular biomarkers for early diagnosis, prevention and personalized therapy, are critically important and highly demanded.

In recent years, number of biomarkers found to be associated with changes in neuron structure and function in schizophrenia patients [11]. With the advancement of the new generation sequencing (NGS) technology and bioinformatics techniques, the ability of humans to understand diseases from the root has greatly upgraded, and more and more disease-related risk genes have been identified. Many NGS studies have shown that mRNAs and the protein it encodes play essential roles in the pathogenesis of schizophrenia. They influence disease manifestation, advancement, and prognosis through their interactions and regulation of signaling pathways. For example, investigation have shown that GLT8D1 and CSNK2B [12], PPP3CC [13], DTNBP1 [14], CSMD1, C10orf26, CACNA1C and TCF4 [15], and ZNF804A [16] expression are altered in schizophrenia patients, it can be used as a biomarkers for diagnosis of schizophrenia. Interestingly, signaling pathways include Akt signaling pathway [17], Wnt signaling pathway [18], MAPK-and cAMP-associated signaling pathways [19], NF-κB signaling pathway [20] and PI3K signaling pathway [21] were observed in schizophrenia. However, our ability to understand the molecular basis of schizophrenia remains limited. In this regard, it is necessary to address the association of genes and signaling pathways in candidate genomes with schizophrenia development.

In this investigation, we screened out the differential expressed genes (DEGs), between normal control and schizophrenia patients in both neural progenitor cells and neurons, from NGS data of Gene Expression Omnibus (GEO) [https://www.ncbi.nlm.nih.gov/geo/] [22]. The key pathways and biomarkers were identified from analysis of gene ontology (GO), REACTOME pathways, protein-protein interaction (PPI) networks, modules, miRNA-hub gene regulatory network and TF-hub gene regulatory network and receiver operating characteristic (ROC) curve analysis tools that offer a new clue to uncover the molecular pathogenesis of schizophrenia.

## Materials and Methods

### Next generation sequencing data source

NGS dataset GSE106589 [23] was downloaded from GEO. GSE106589 containing 46 cases of schizophrenia and 33 normal control cases were obtained from the GPL16791 Illumina HiSeq 2500 (Homo sapiens) platform. All cases were of human source.

### Identification of DEGs

The differential expression analysis on mRNA was performed using the DESeq2 package [24] in R software. The DEGs between schizophrenia and normal control were identified using a P-value of <0.05 and |fold change| > 0.69 for up regulated genes and |fold change| < -0.51 for down regulated genes as the cutoff for screening. The volcano map and heatmap of the DEGs were respectively generated using the ggplot2 and gplot packages in R software.

### GO and pathway enrichment analyses of DEGs

GO functional annotation (http://www.geneontology.org) [25] includes biological processes (BP), cellular component (CC), and molecular function (MF), which can be used to clarify the potential biological functions of the enriched genes. Pathway enrichment analysis can be used to identify the main biochemical metabolic pathways and signal transduction pathways involved in enriched genes. GO and REACTOME (https://reactome.org/) [26] pathway enrichment analysis of the DEGs were performed using the g:Profiler (http://biit.cs.ut.ee/gprofiler/) [27]. A P-value of ≤ 0.05 was used as the cutoff for screening.

### Construction of the PPI network and module analysis

The International Molecular Exchange Consortium (IMEx) (https://www.imexconsortium.org/) [28], the most commonly used online tool for PPI network analysis in the biomedical field, was used to develop the PPI network of DEGs. Finally, the PPI network was visualized by using the Cytoscape software (V3.10.1; http://cytoscape.org/) [29]. The Network Analyzer plug-in was used to calculate node degree [30], betweenness [31], stress [32] and closeness [33] of hub genes in the PPI network. The PEWCC [34] of Cytoscape was carried out to module analyze and visualize the result of PPI network and the key modules with the highest score were selected for visualization display.

### Construction of the miRNA-hub gene regulatory network

miRNA-hub gene was predicted by the miRNet database (https://www.mirnet.ca/) [35]. The miRNAs of interaction in databases (TarBase, miRTarBase, miRecords, miRanda (S mansoni only), miR2Disease, HMDD, PhenomiR, SM2miR, PharmacomiR, EpimiR, starBase, TransmiR, ADmiRE and TAM 2) were as the predicted miRNA that might regulate schizophrenia. The miRNA- hub gene network was visualized by cytoscape software [29].

### Construction of the TF-hub gene regulatory network

TF-hub gene was predicted by the NetworkAnalyst database (https://www.networkanalyst.ca/) [36]. The TFs of interaction in database (Jasper) were as the predicted TF that might regulate schizophrenia. The TF- hub gene network was visualized by cytoscape software [29].

### Receiver operating characteristic curve (ROC) analysis

ROC curve analysis, which yields indicators of certainty such as the area under the curve (AUC), provides the crucial principle and explanation for distinguishing between the specificity and sensitivity of diagnostic performance of hub genes. We used the pROC package of R software [37] to conduct our ROC curve analysis.

## Results

### Identification of DEGs

The NGS dataset GSE106589 was downloaded from the GEO database, and 955 schizophrenia related DEGs were obtained by a differential analysis, of which 478 were highly expressed (p < 0.05, |fold change| > 0.69) and 477 were poorly expressed ((p < 0.05, |fold change| < -0.51) (Table 1). To visualize the DEGs, we constructed volcano plot (Fig.1) and heatmap (Fig.2).

**Fig. 1.**
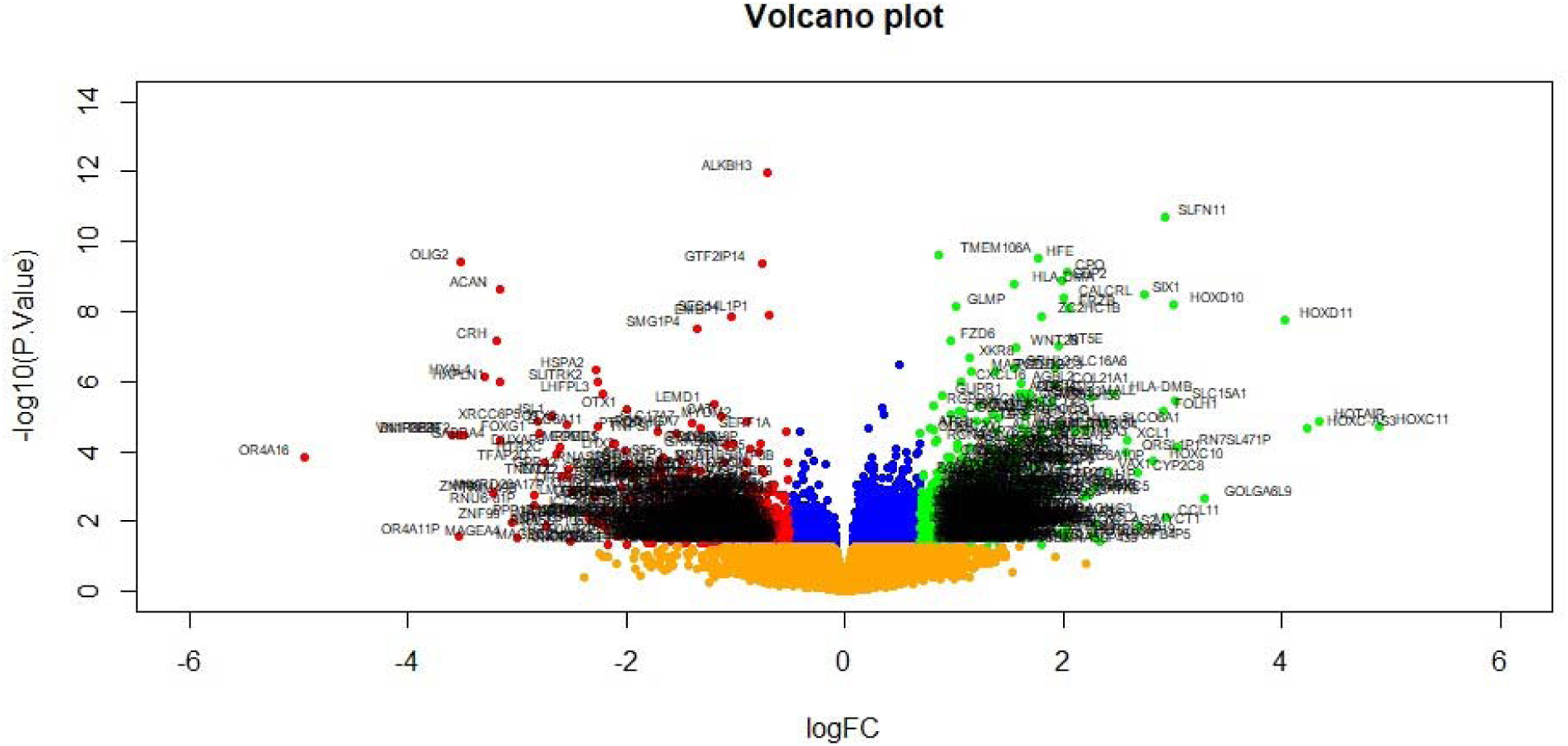
Volcano plot of differentially expressed genes. Genes with a significant change of more than two-fold were selected. Green dot represented up regulated significant genes and red dot represented down regulated significant genes.

**Fig. 2.**
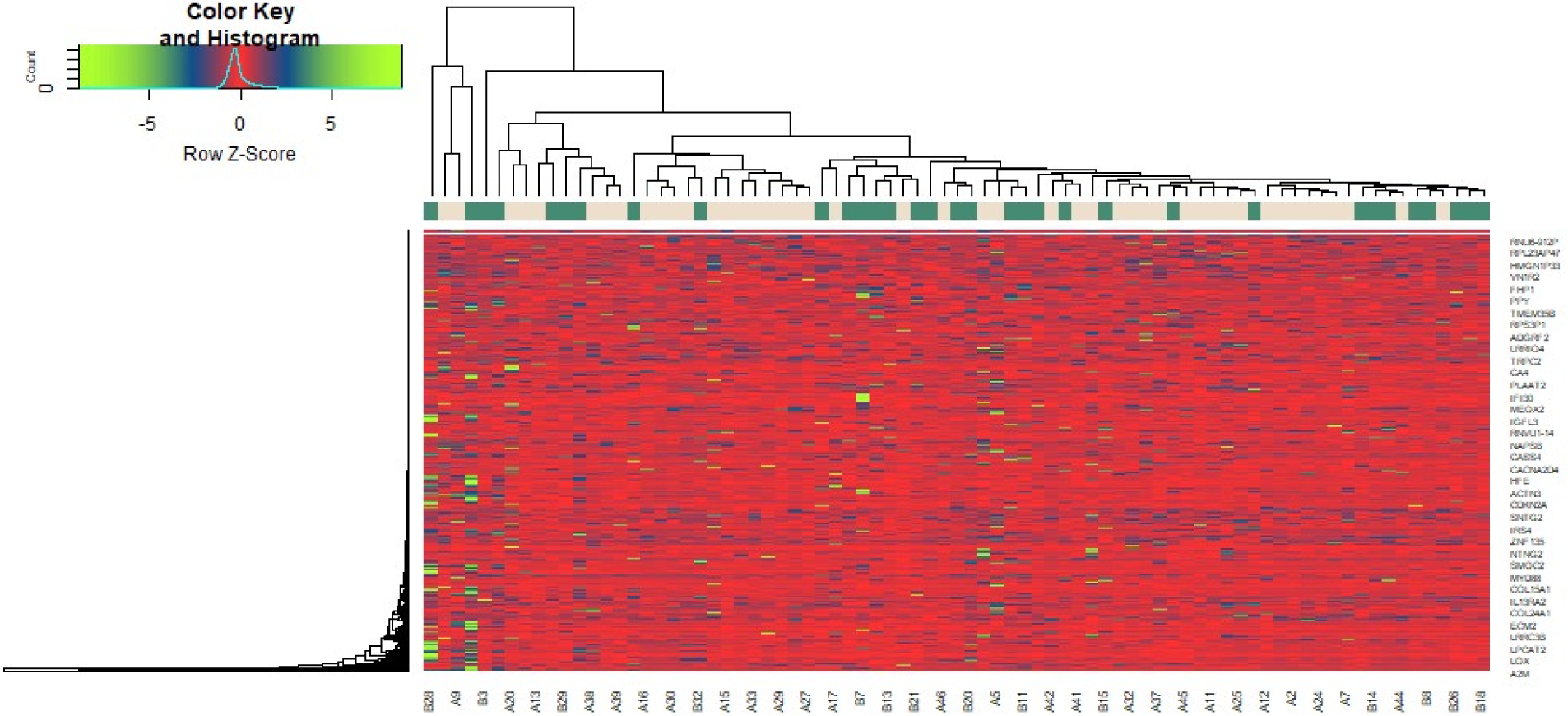
Heat map of differentially expressed genes. Legend on the top left indicate log fold change of genes. (A1 – A46 = schizophrenia samples; B1 – B 33 = Normal control samples)

**Table 1.**
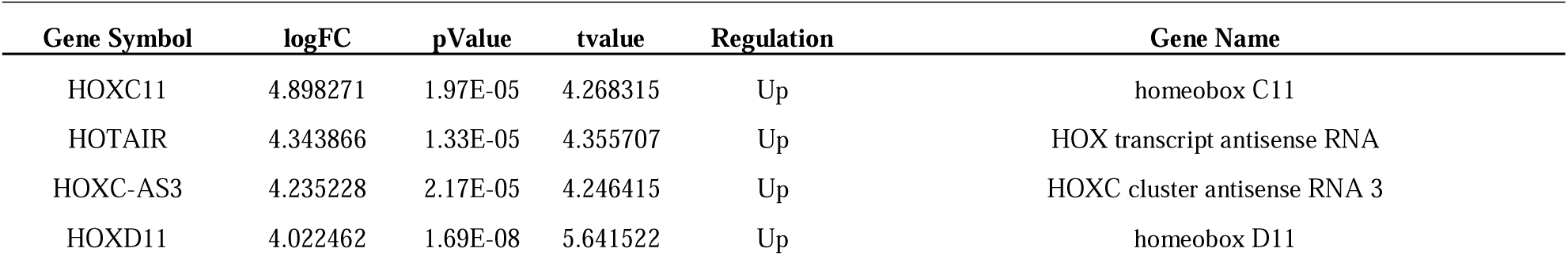
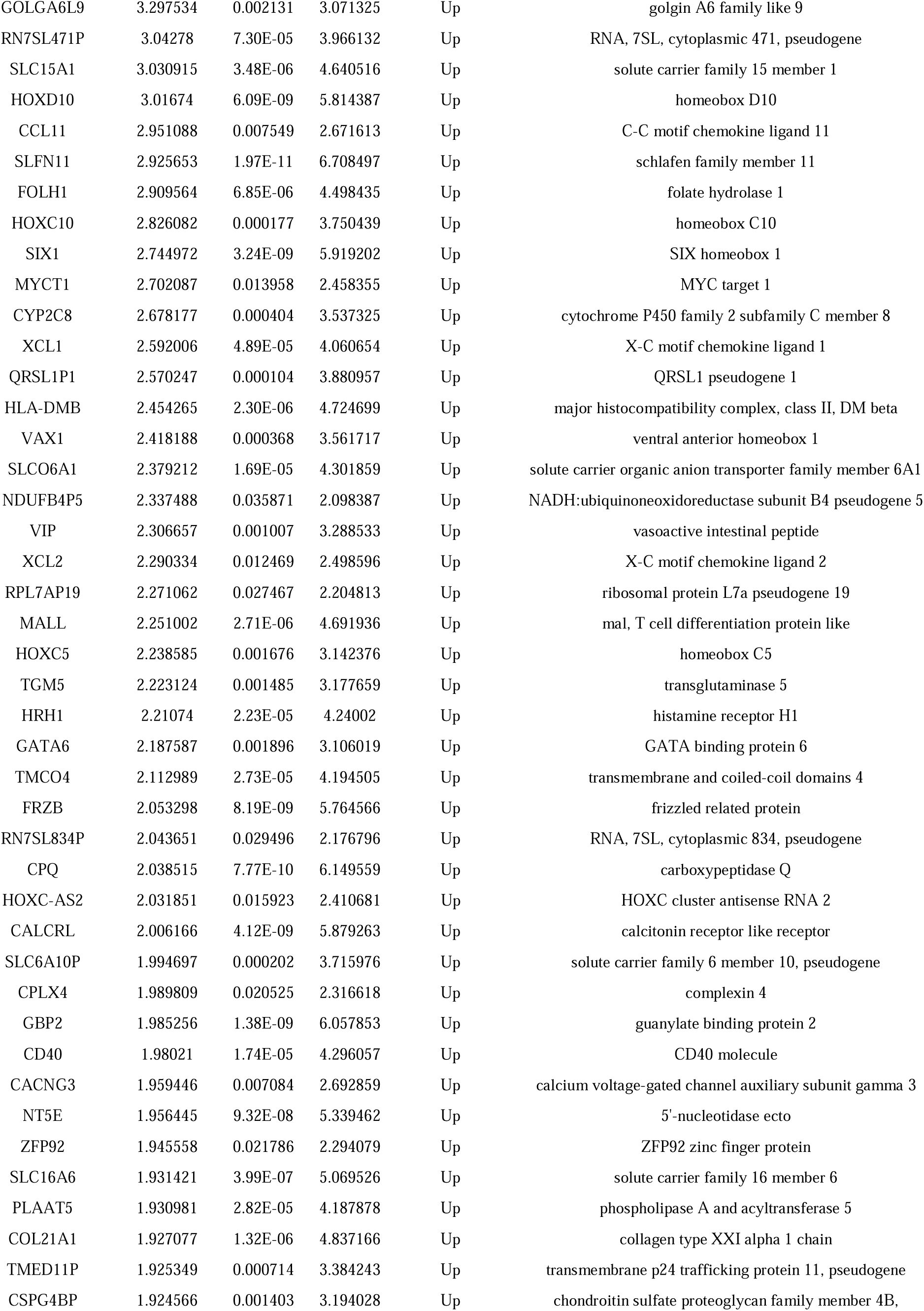
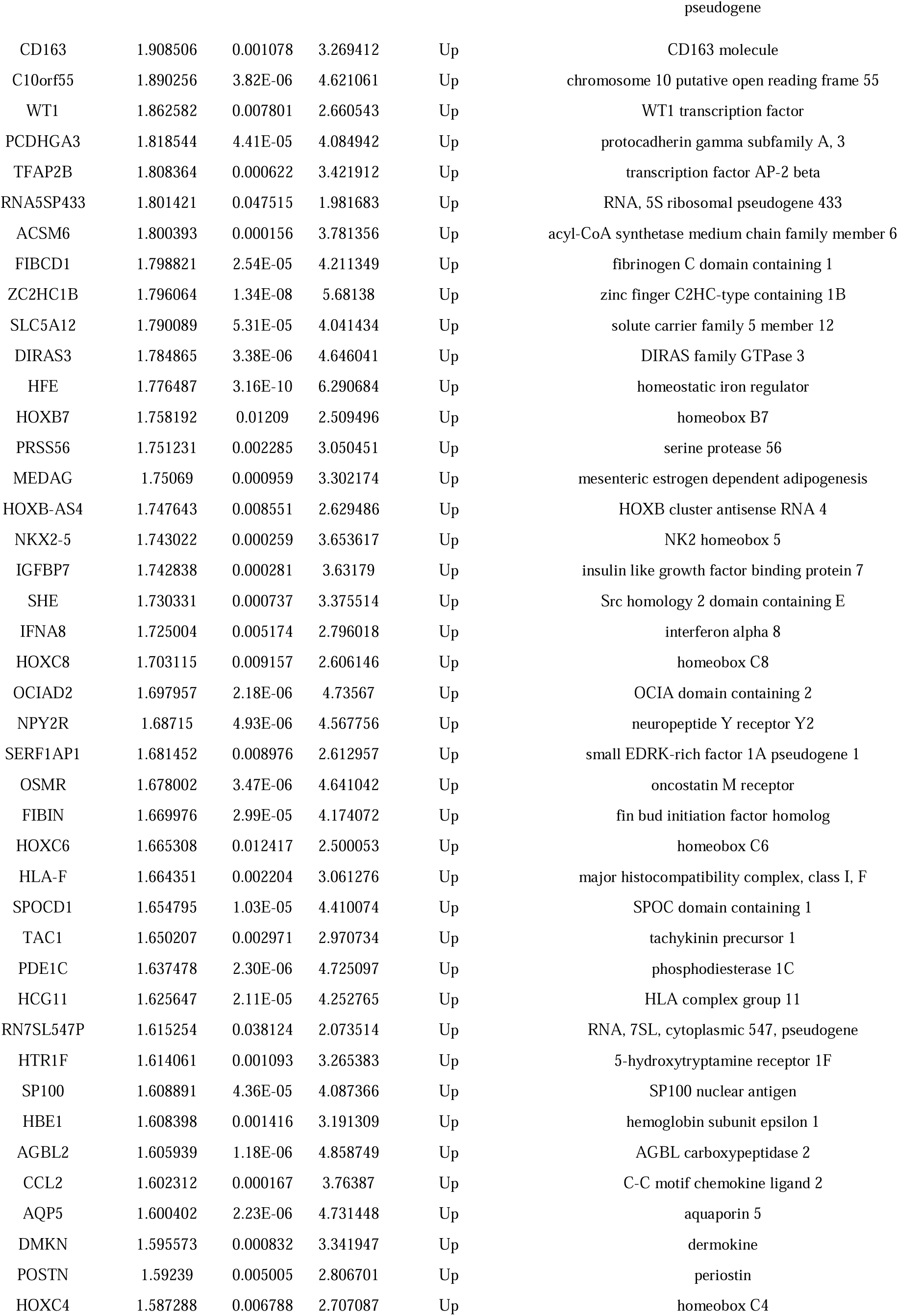
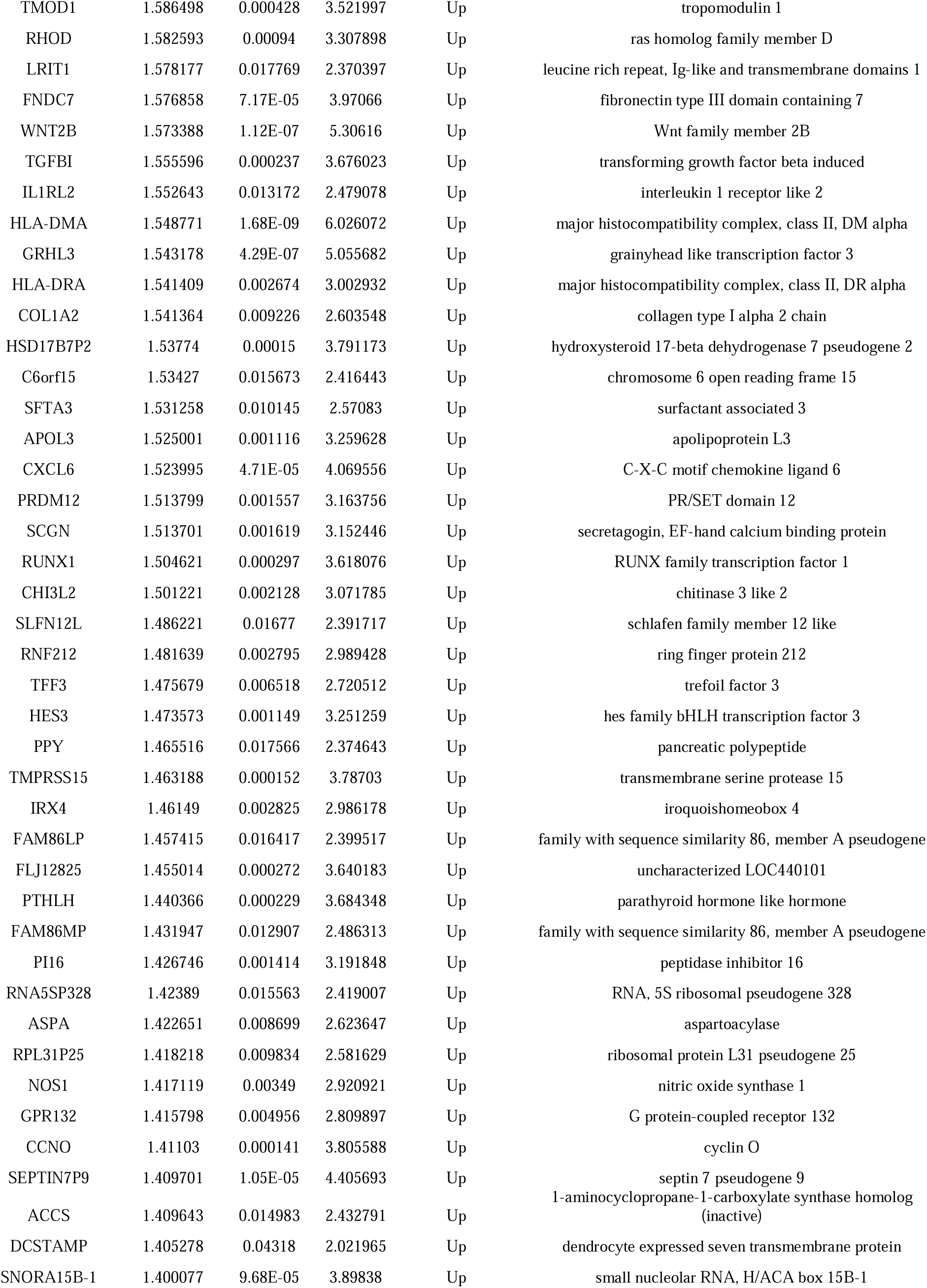
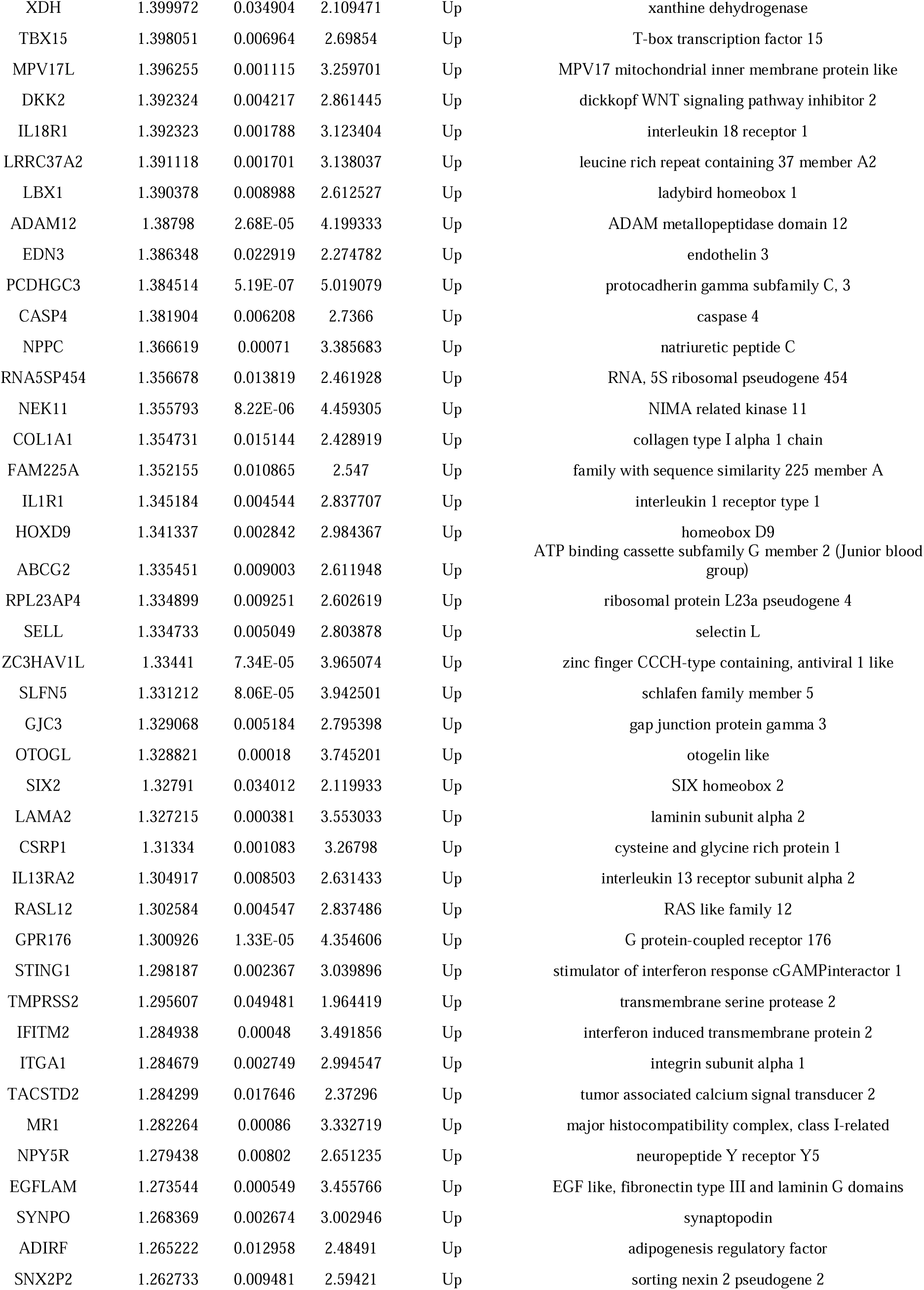
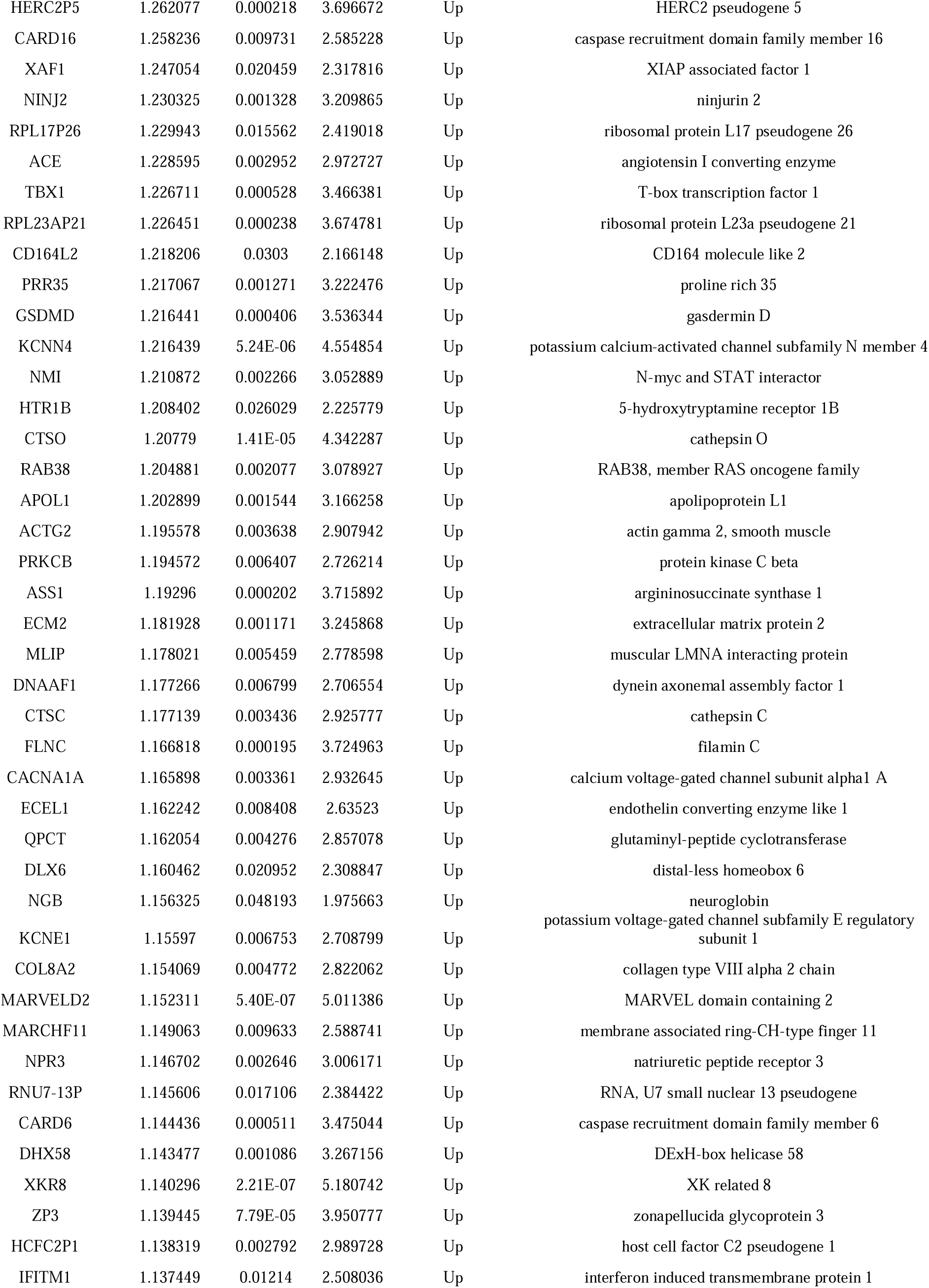
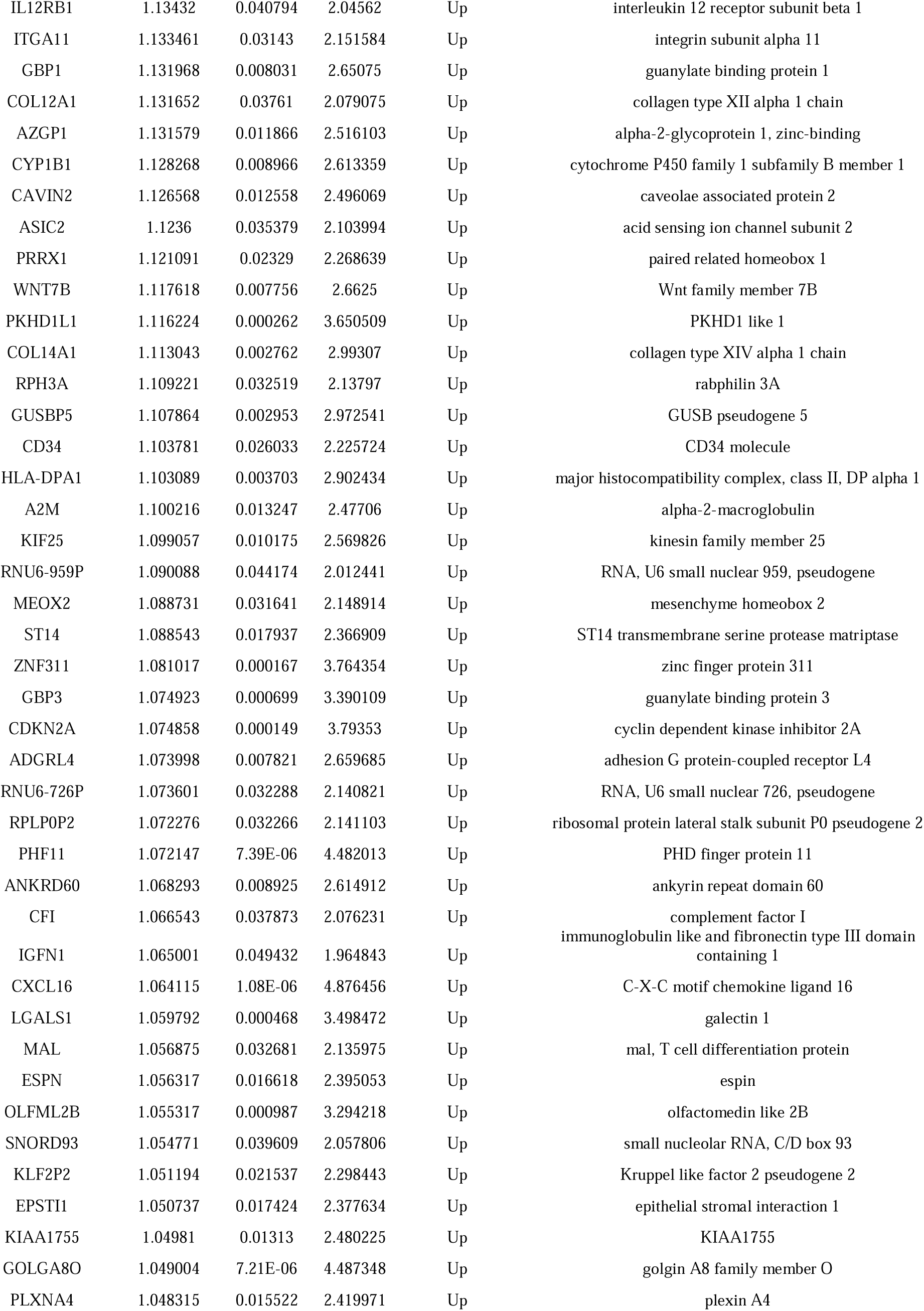
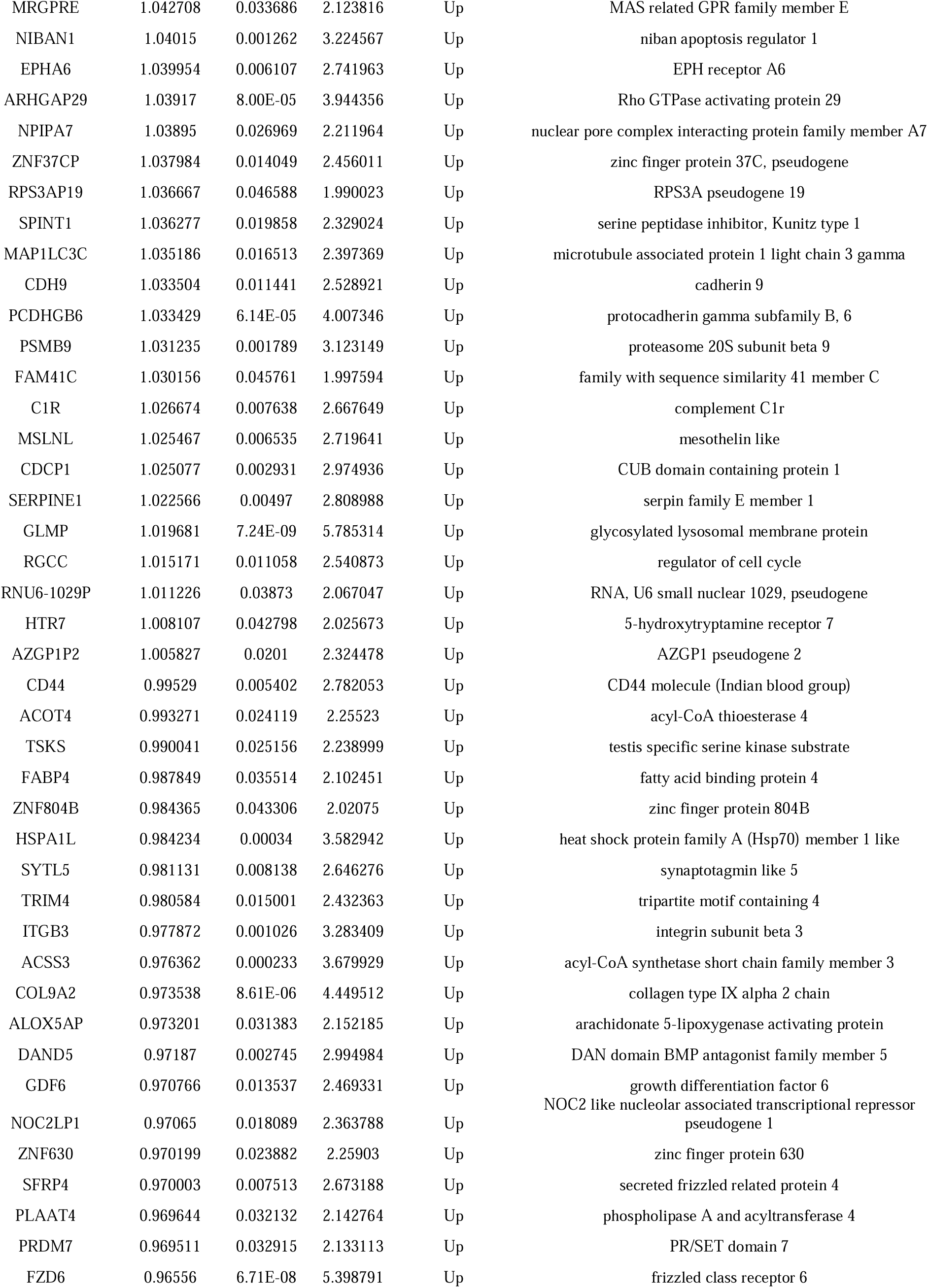
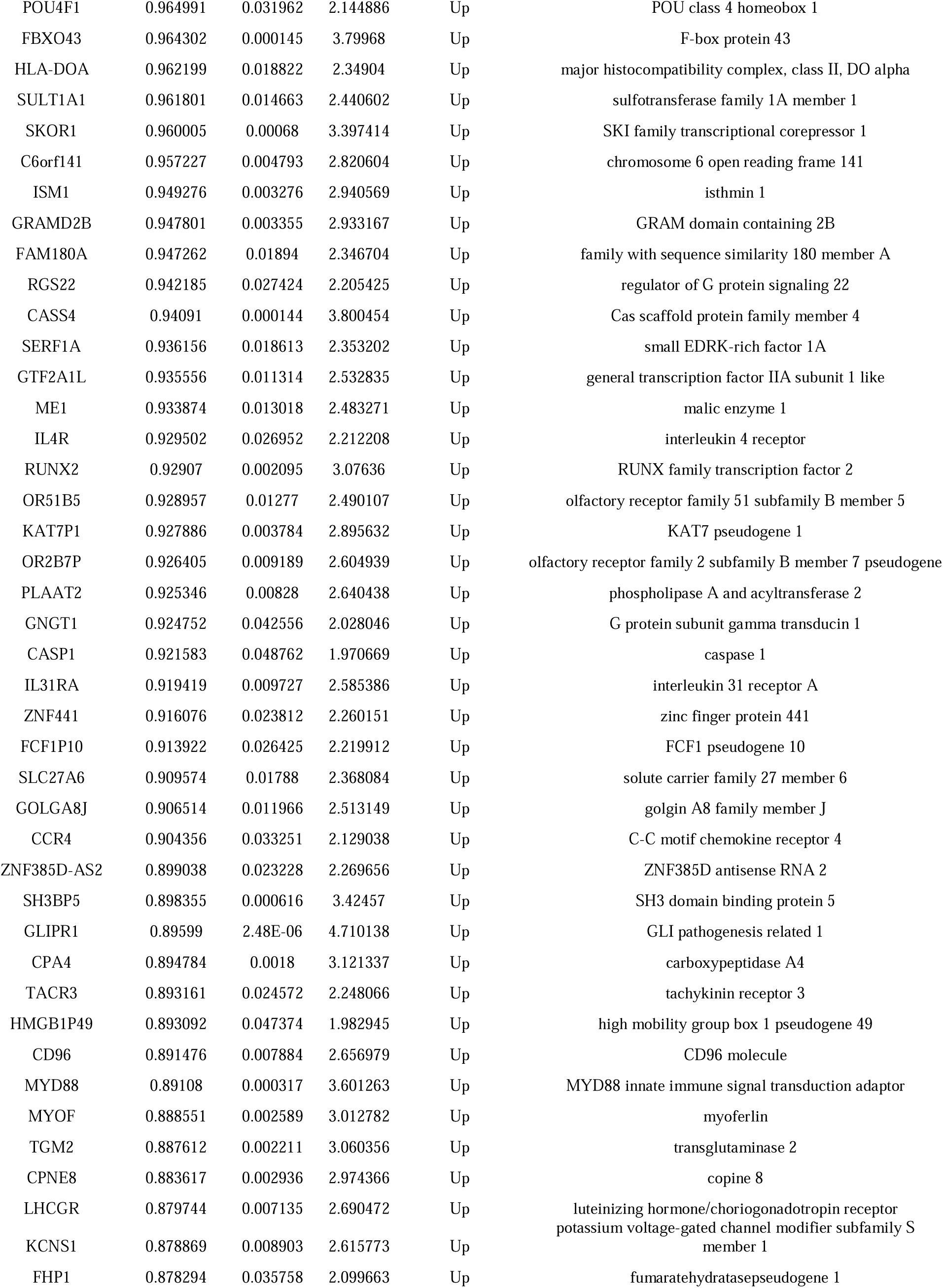
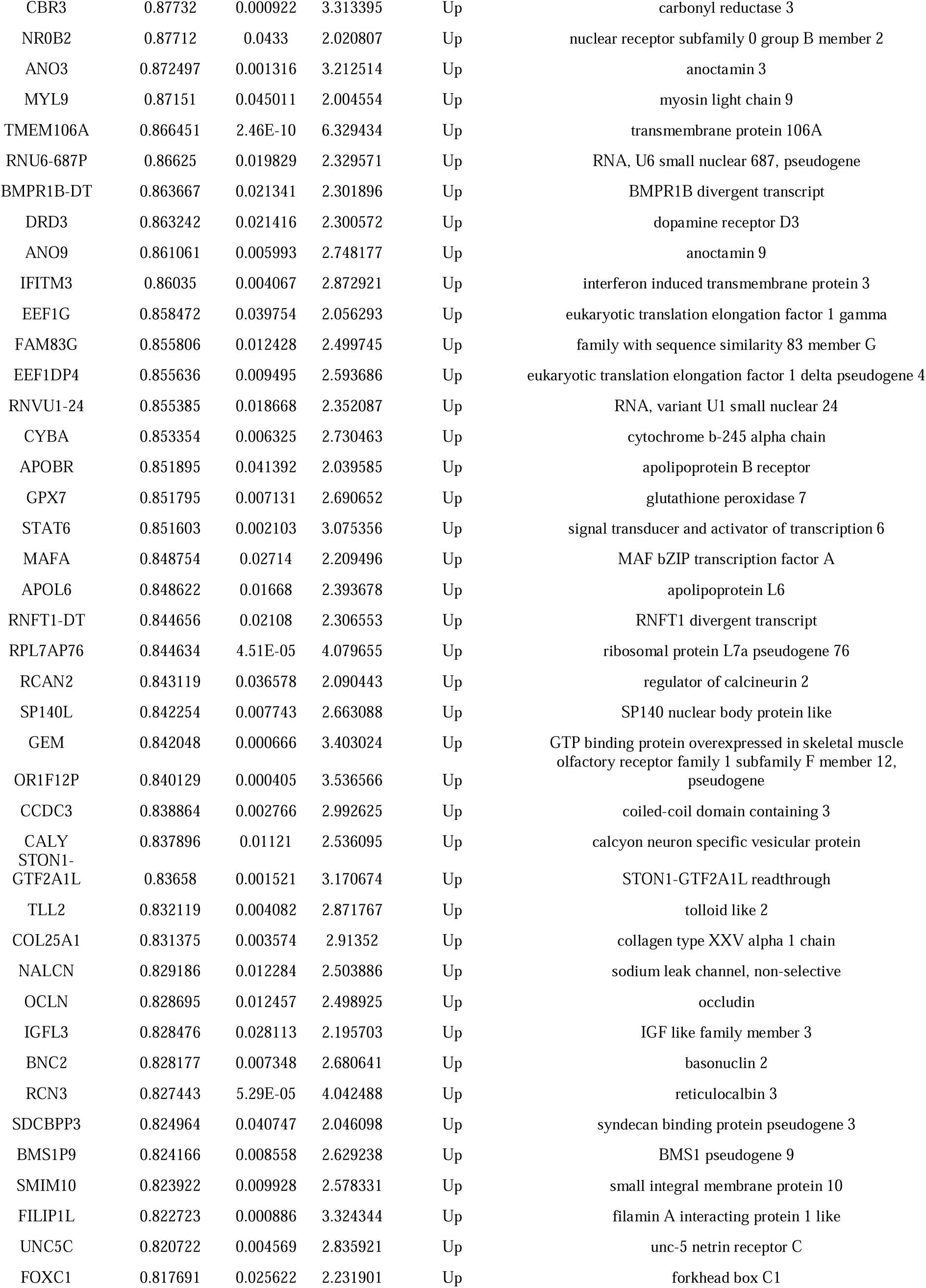
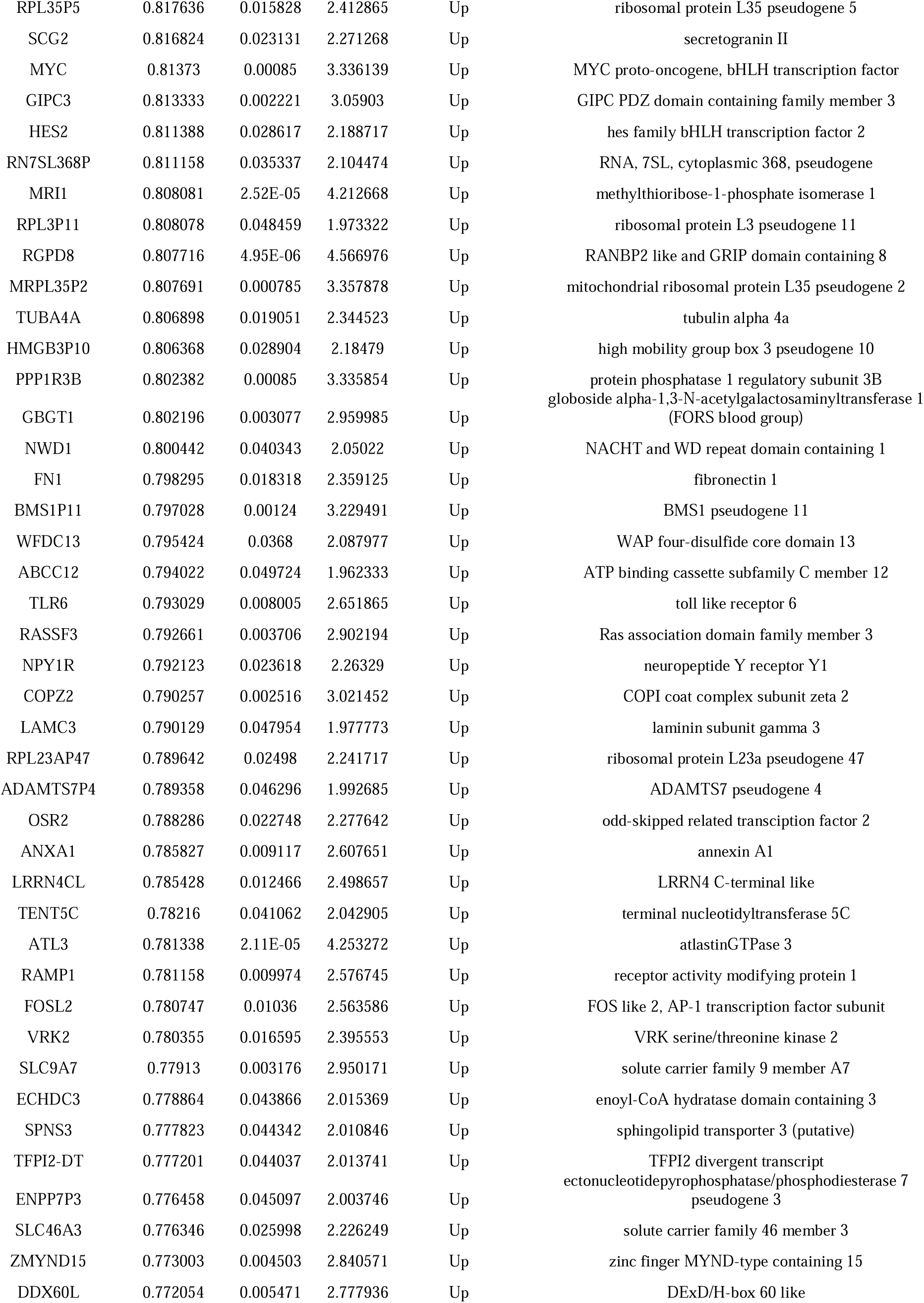
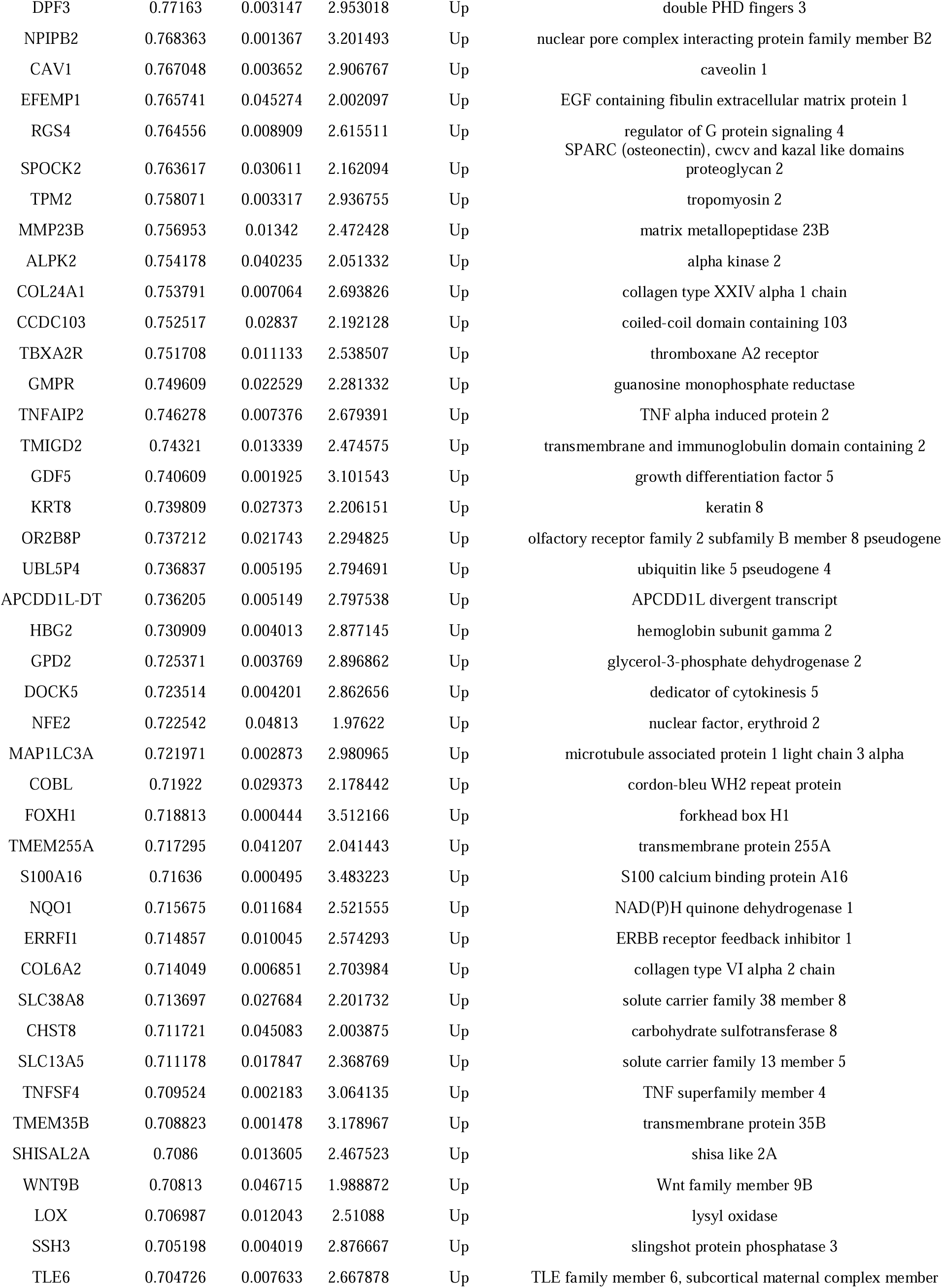
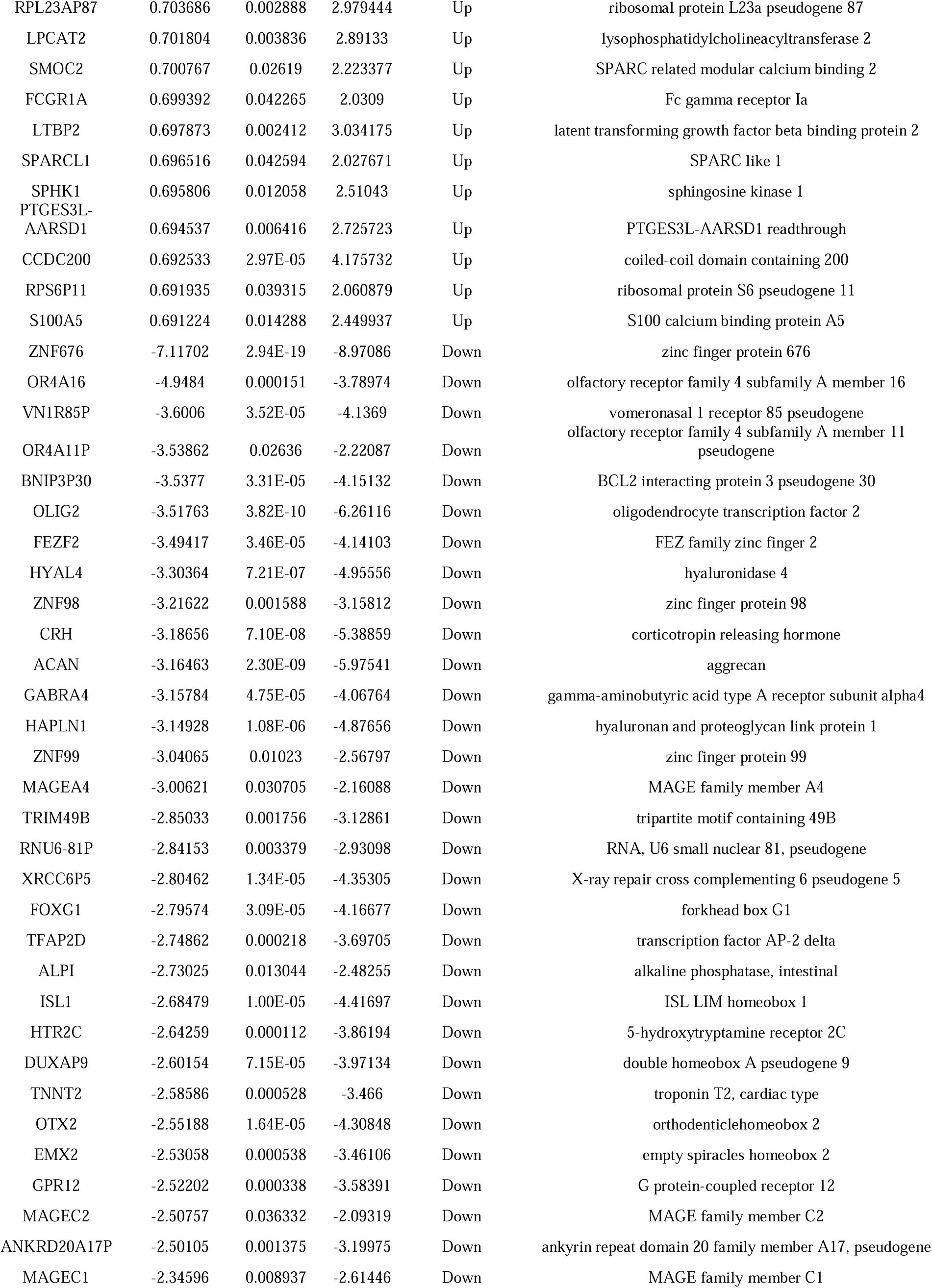
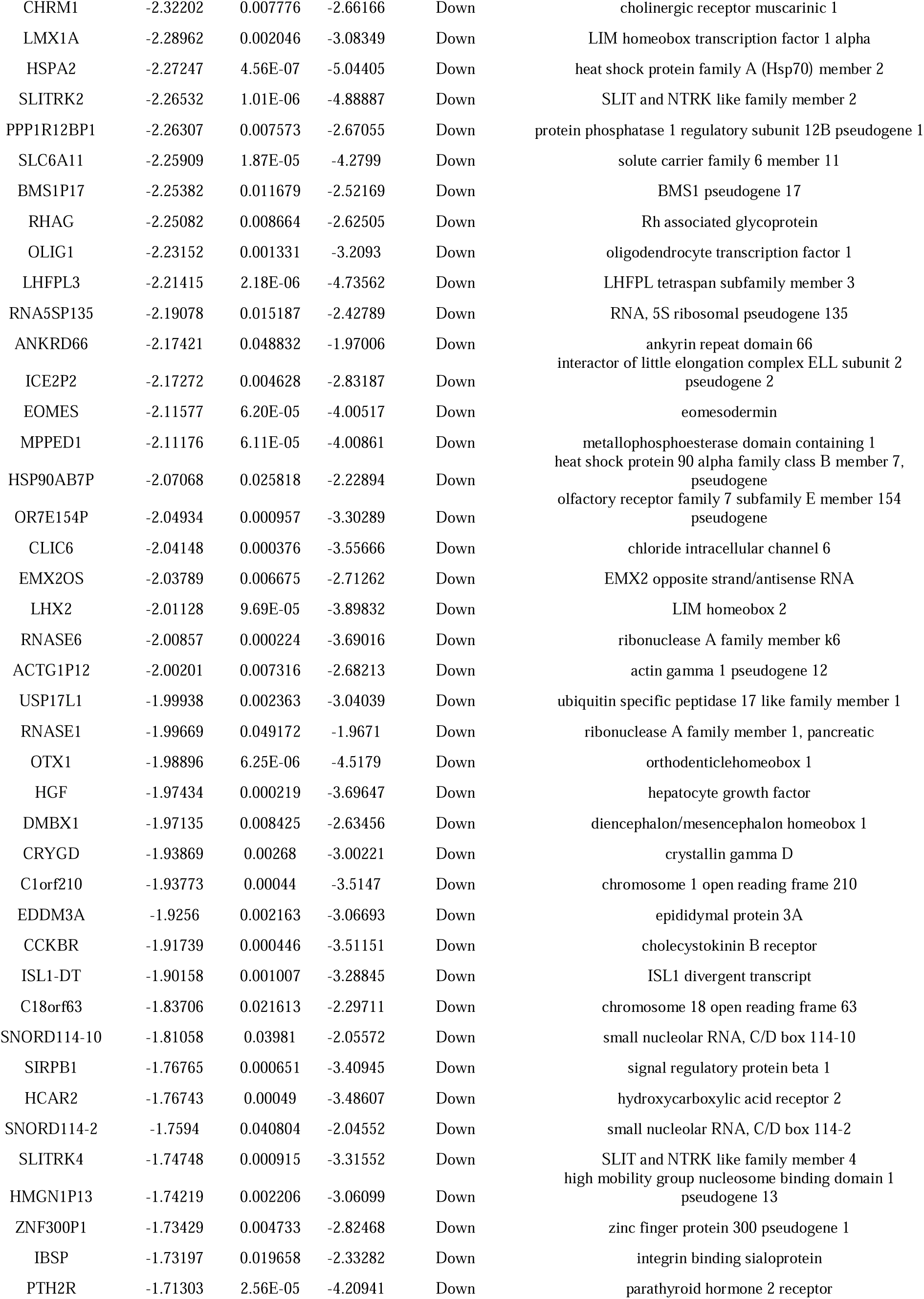
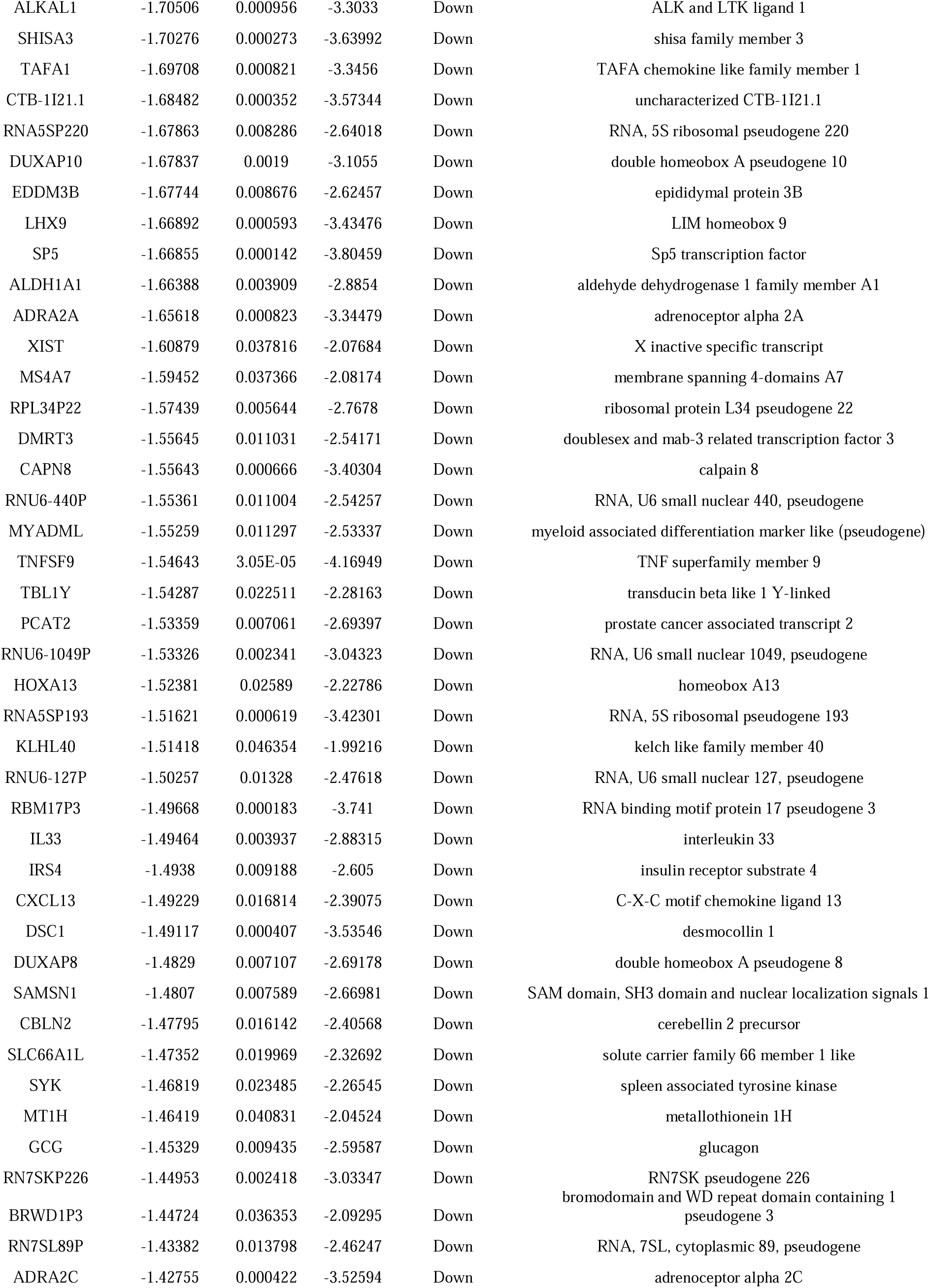
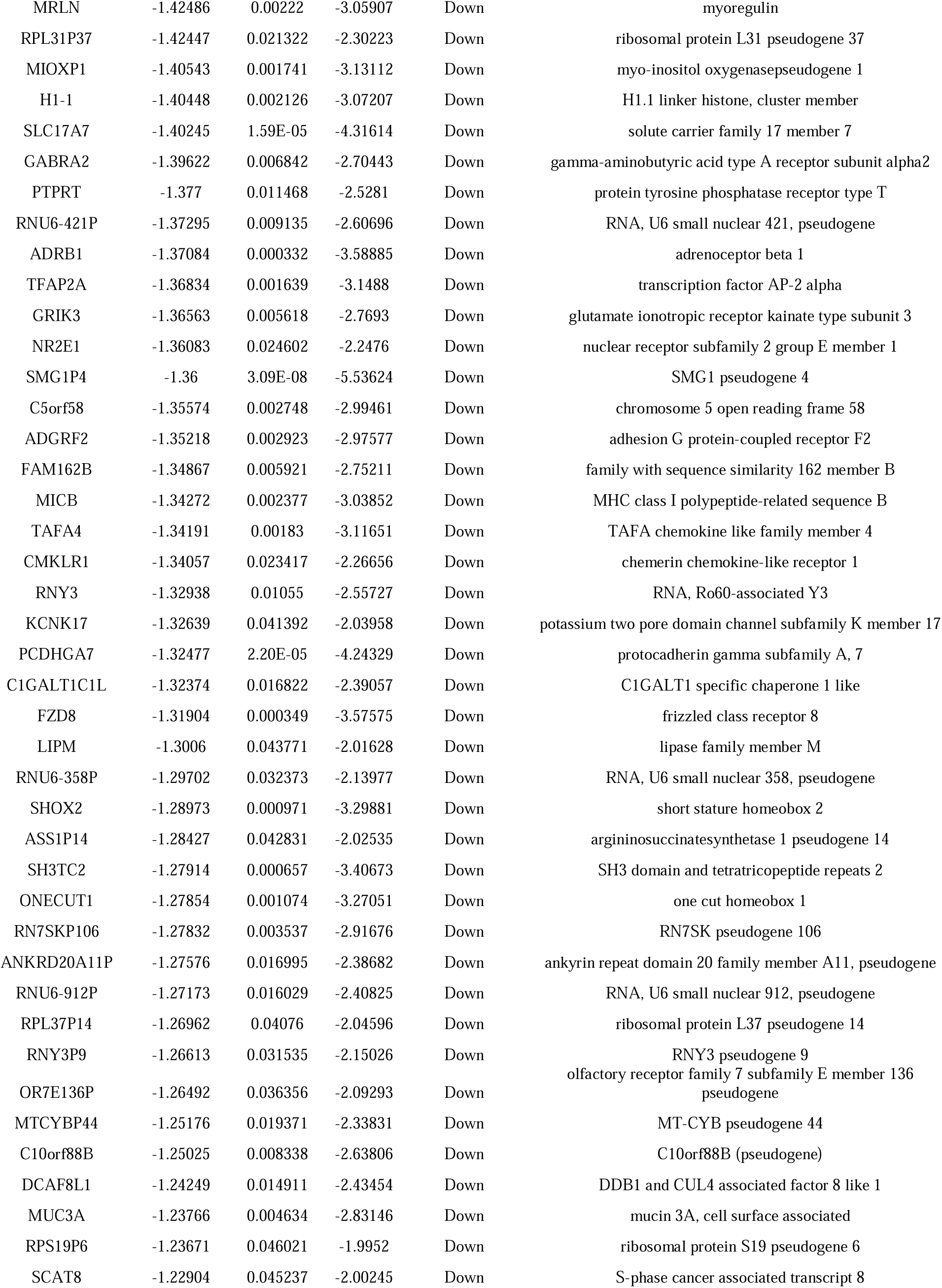
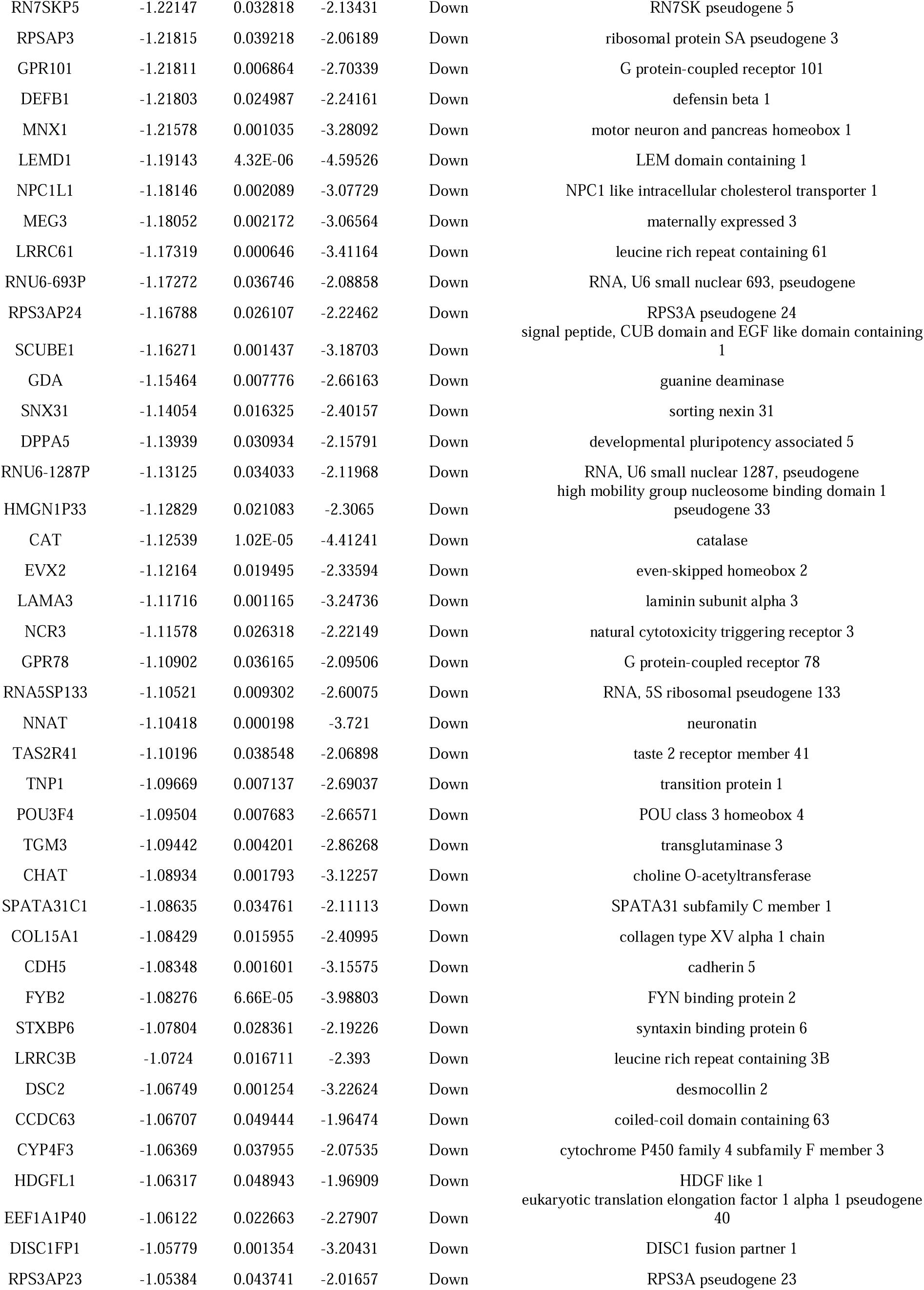
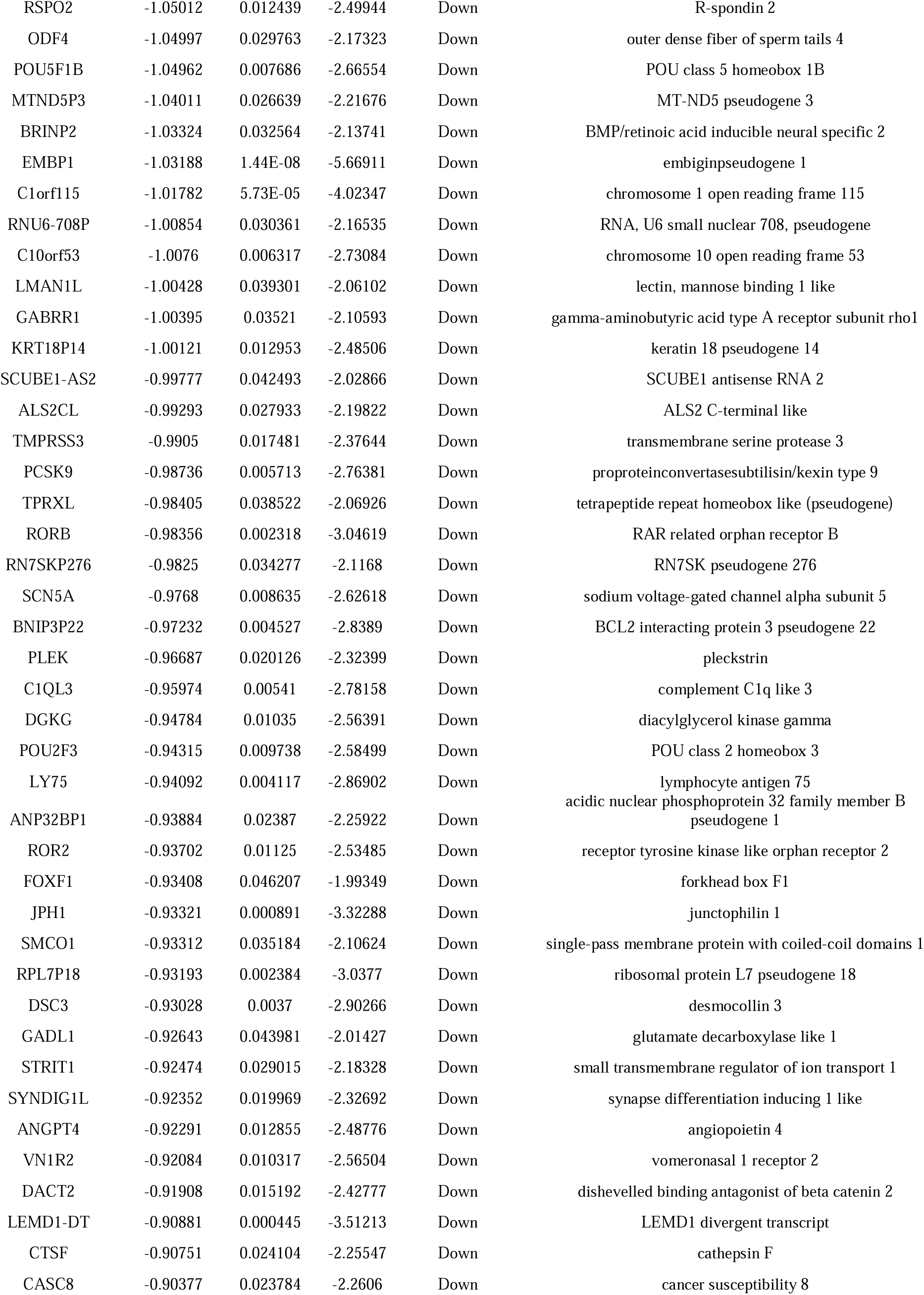
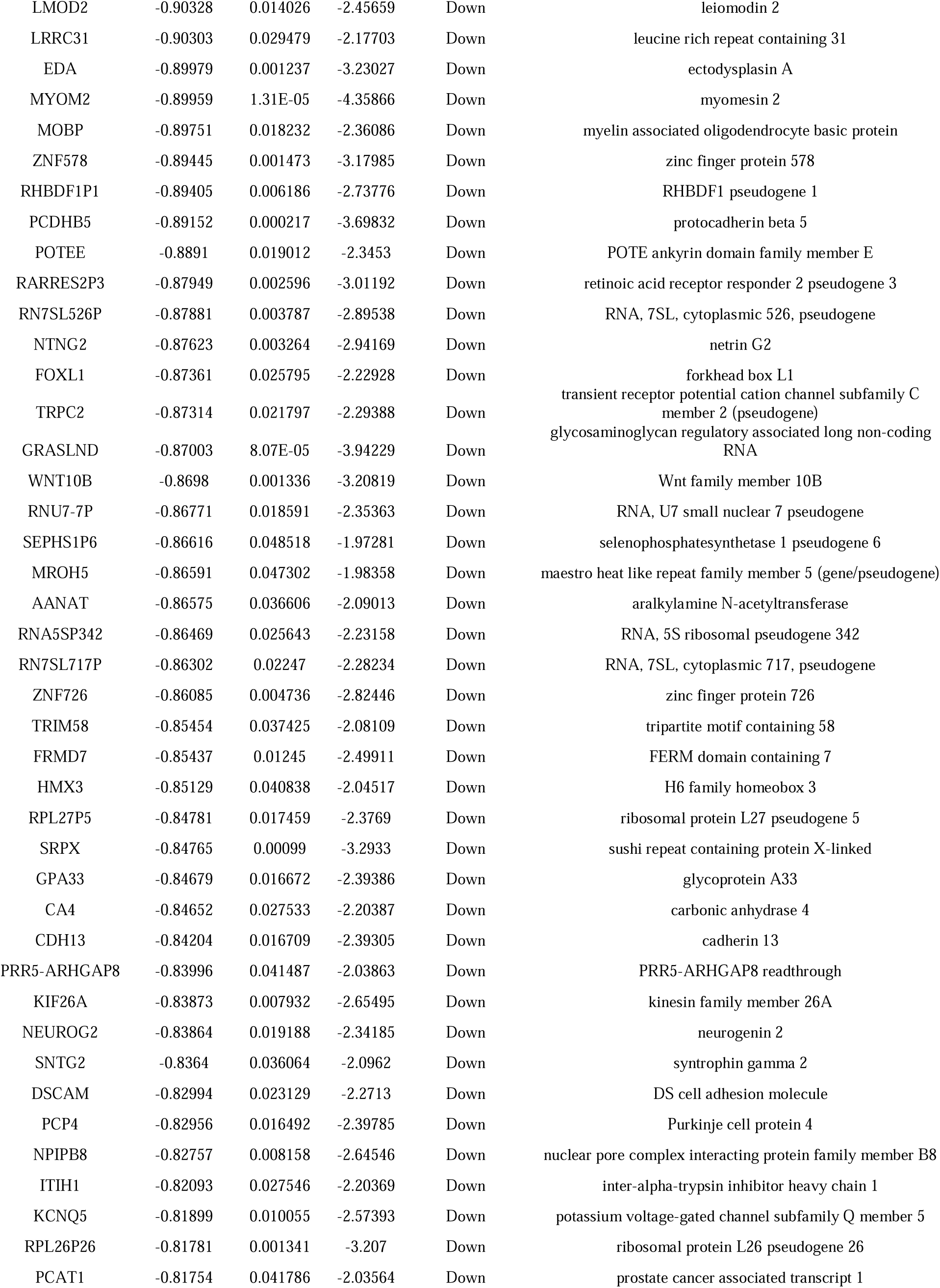
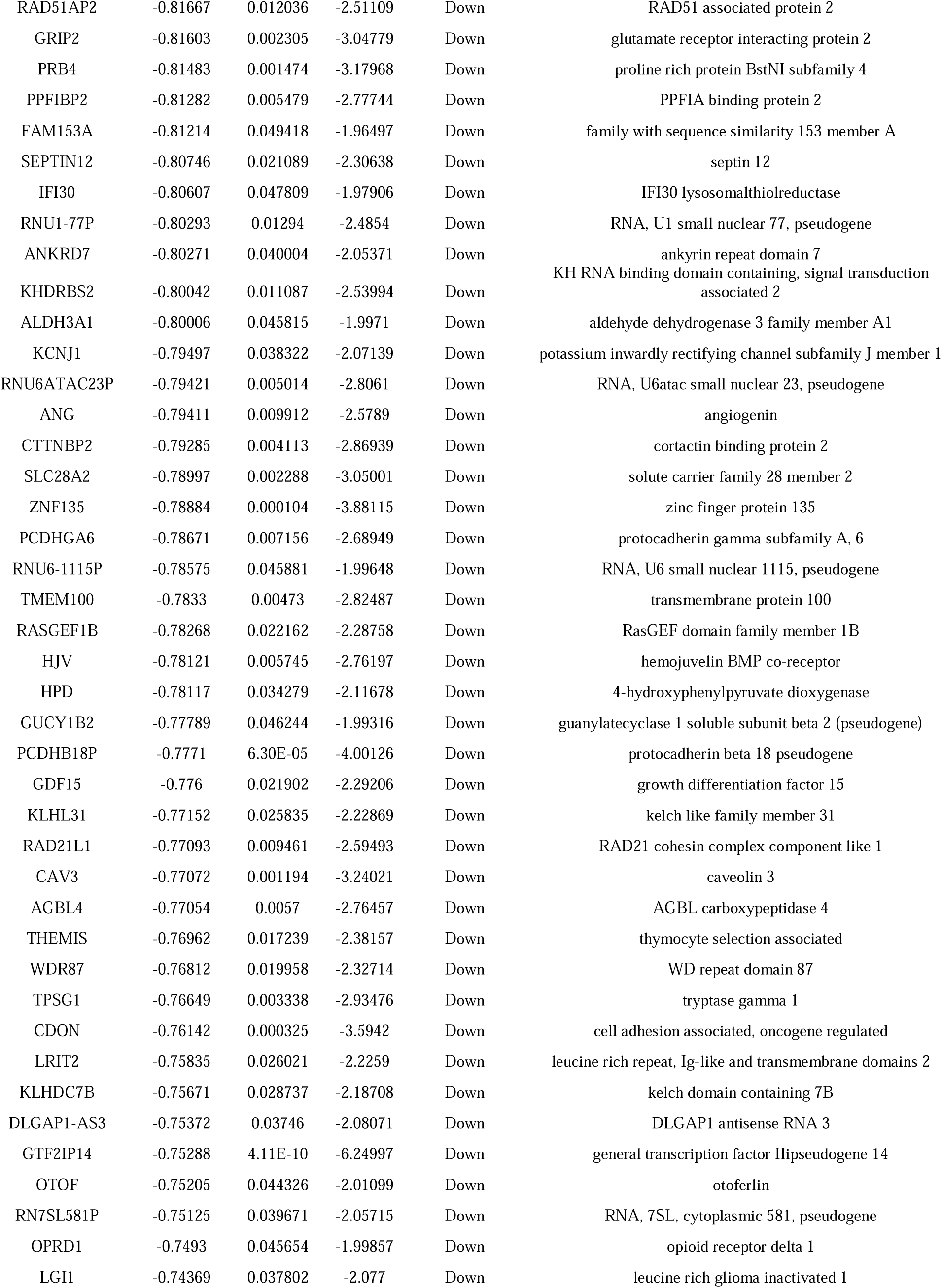
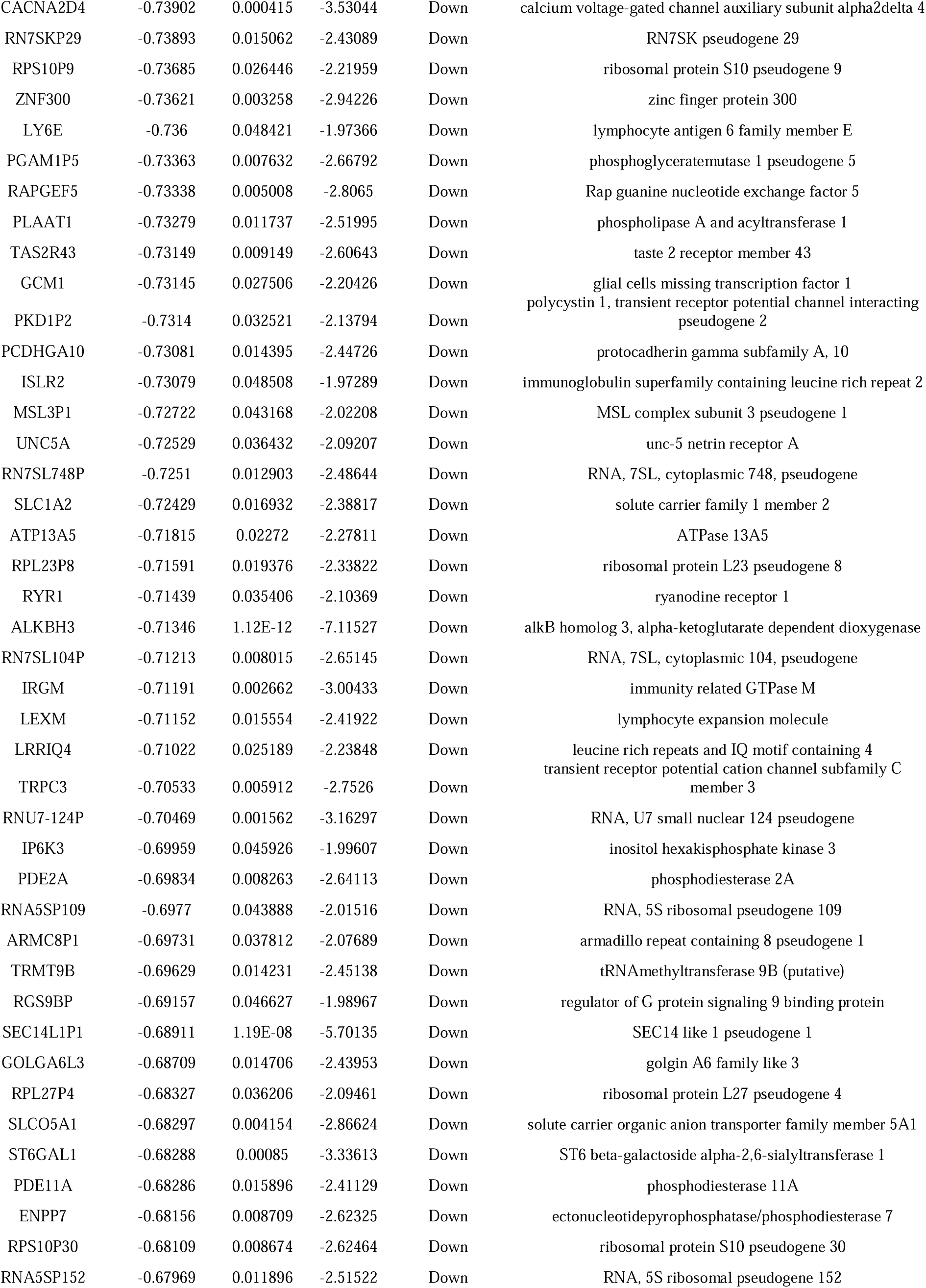
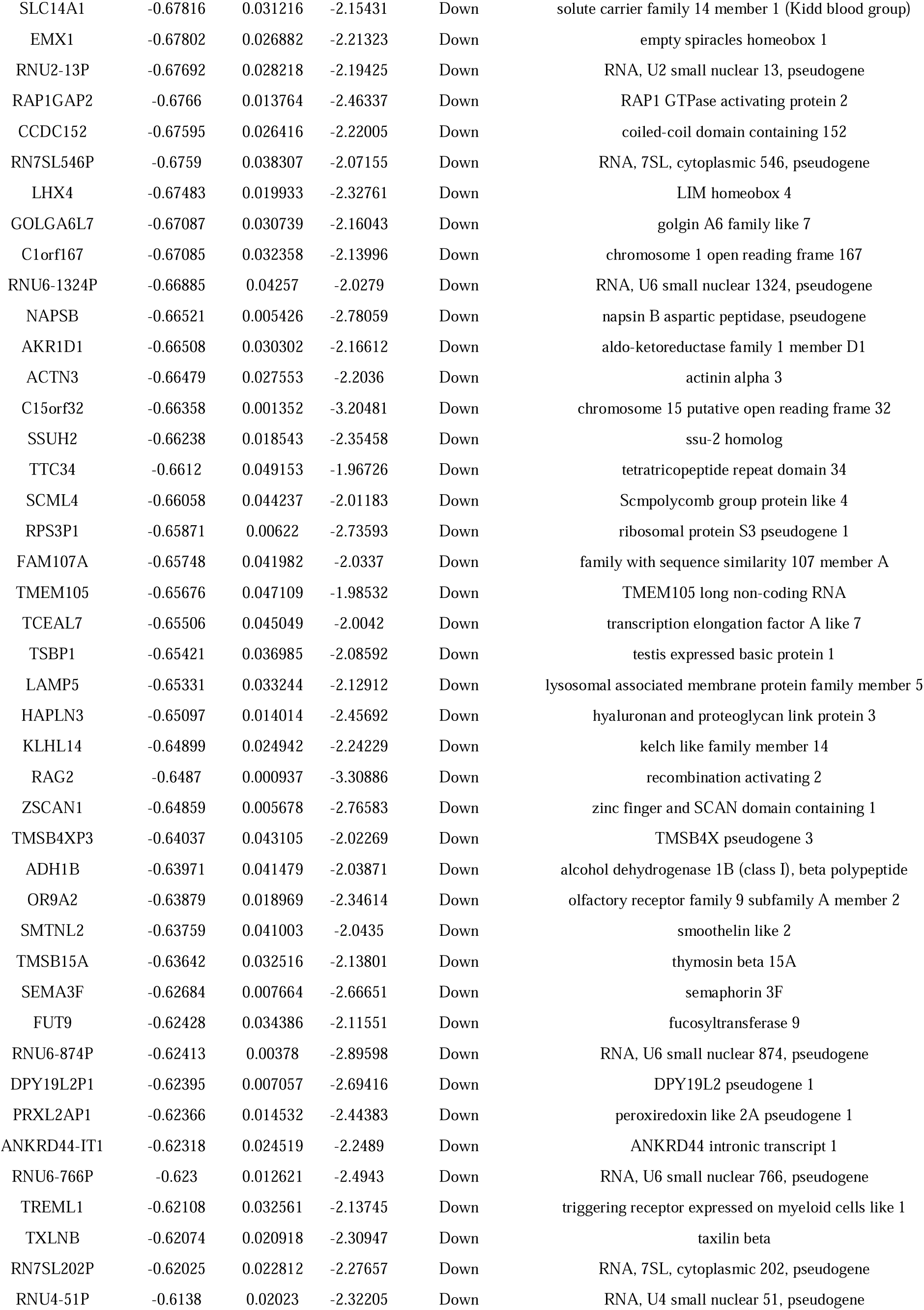
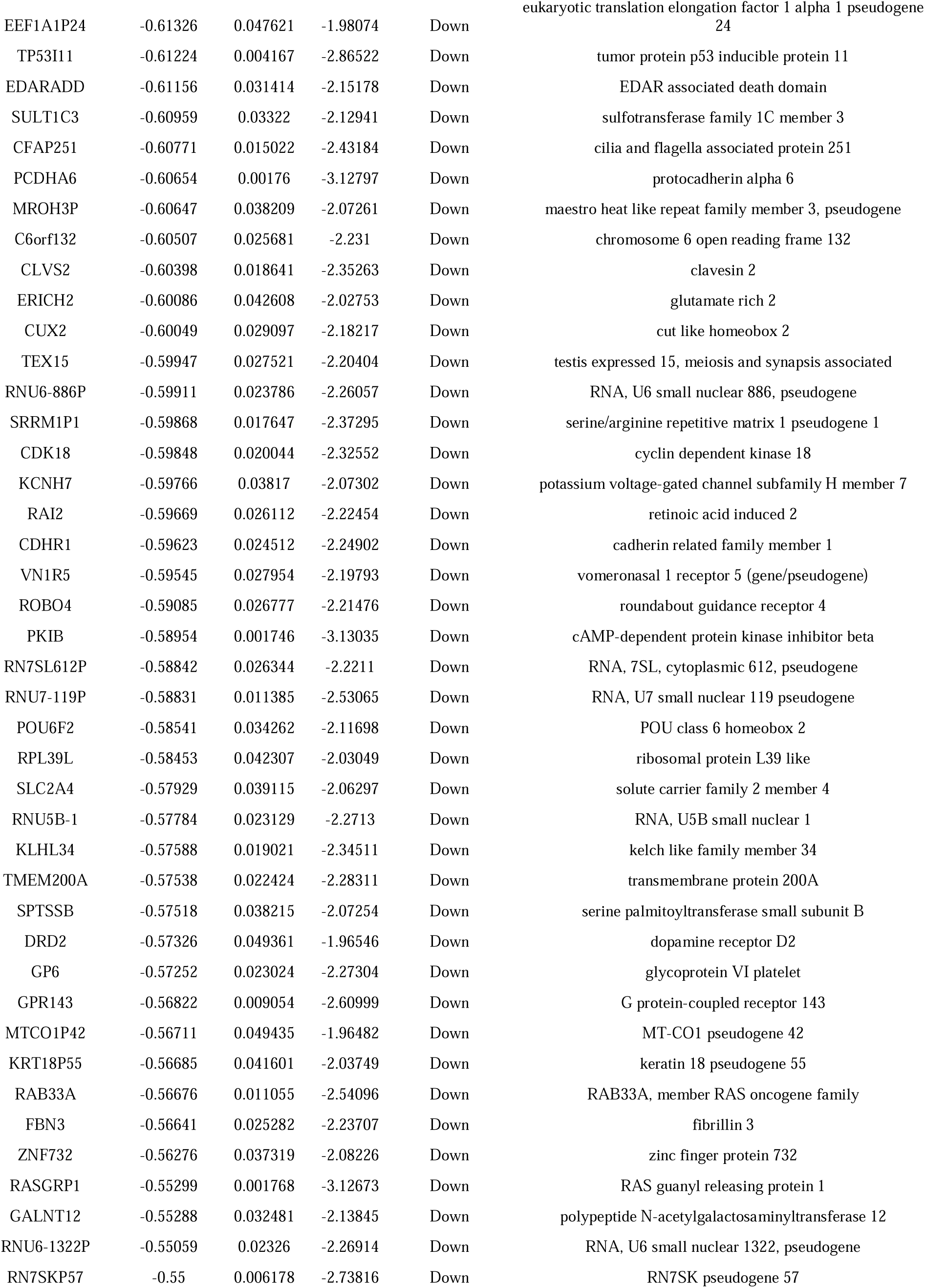
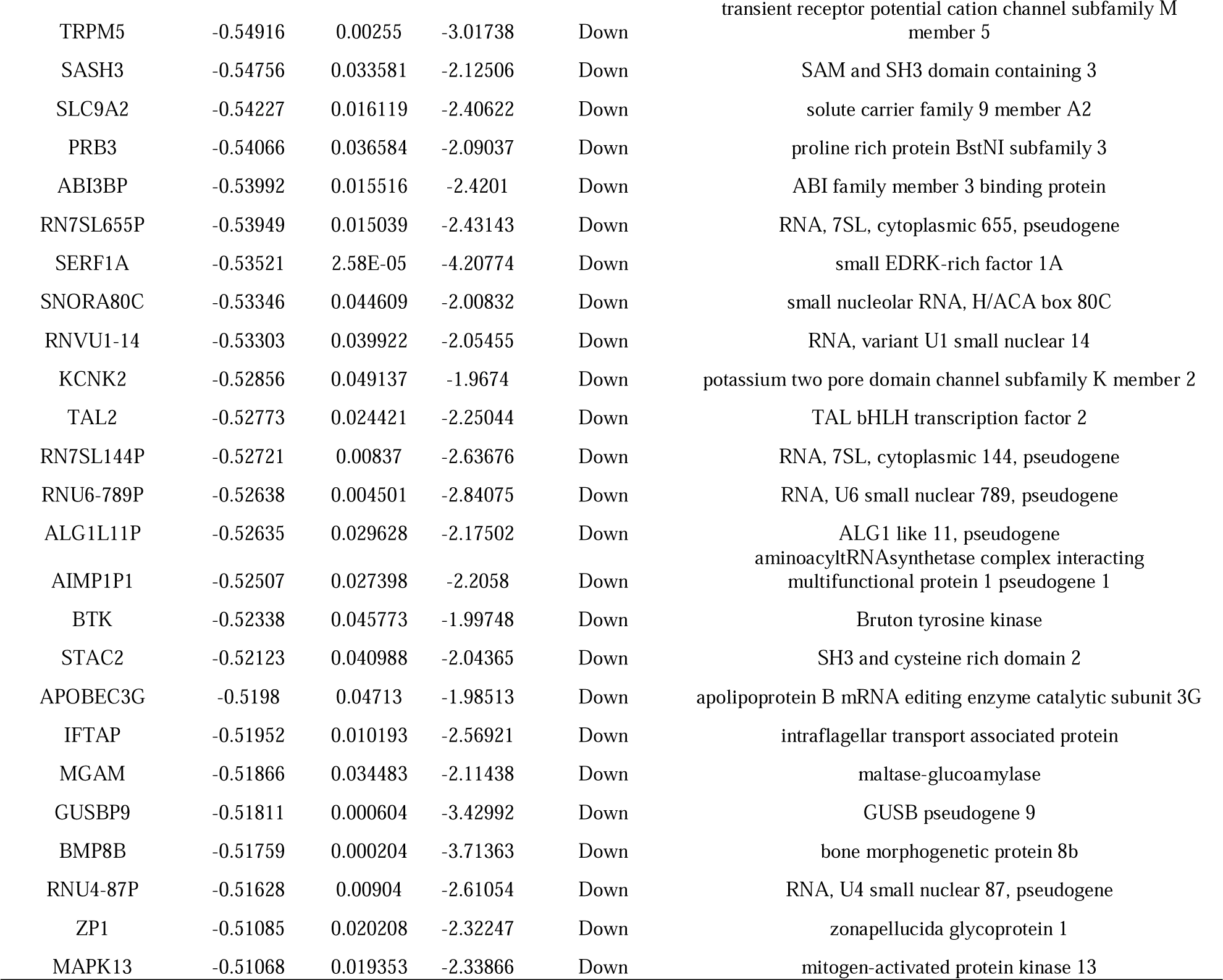
The statistical metrics for key differentially expressed genes (DEGs)

### GO and pathway enrichment analyses of DEGs

GO functional annotation and REACTOME pathway enrichment analysis were performed for the DEGs from the NGS dataset. The significant results are presented in Table 2. In the BP group, the up regulated genes were mainly clustered in multicellular organismal process and developmental process, and the down regulated genes were mainly clustered in regulation of cellular process and biological regulation. For the CC group, the up regulated genes were primarily clustered in cell periphery and cellular anatomical entity. The down regulated genes were primarily clustered in cell periphery and plasma membrane. The up regulated genes in the MF group were mostly clustered in DNA-binding transcription activator activity, RNA polymerase II-specific and signaling receptor binding, and the down regulated genes were mostly clustered in double-stranded DNA binding and sequence-specific DNA binding. The top significantly enriched REACTOME pathways for the DEGs were also displayed by the g:Profiler online software and are presented in Table 3. The up regulated genes were associated with GPCR ligand binding and extracellular matrix organization, while the down regulated genes were involved in amine ligand-binding receptors and signaling by GPCR.

**Table 2.**
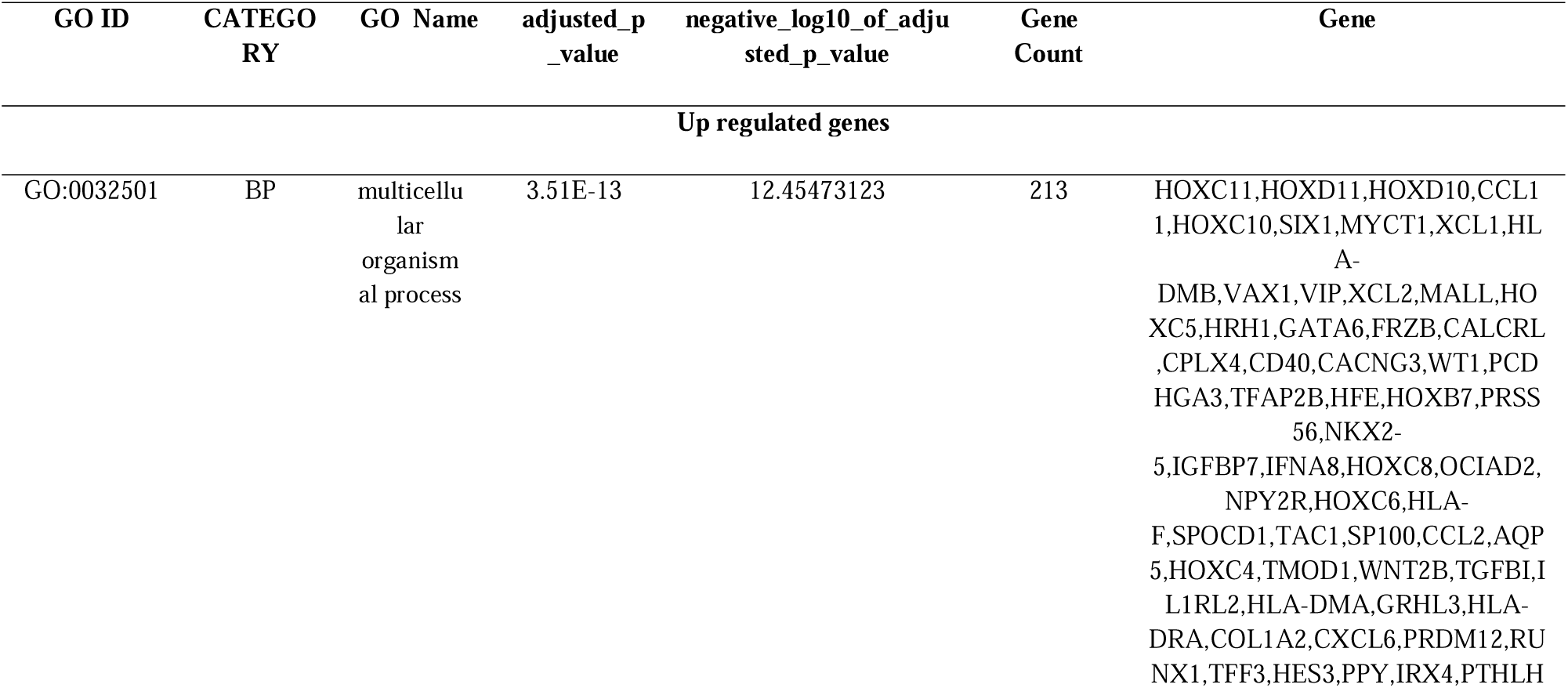
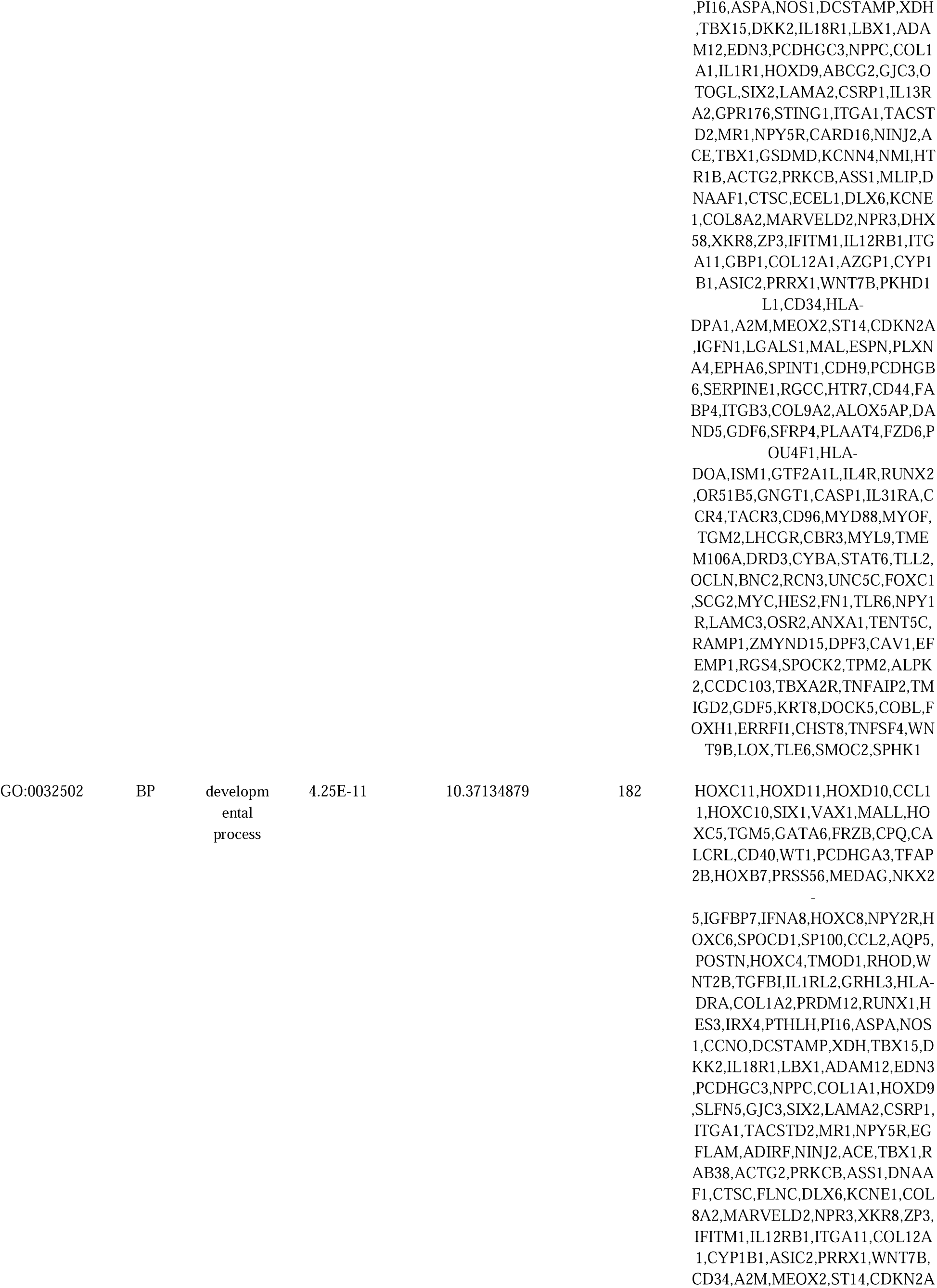
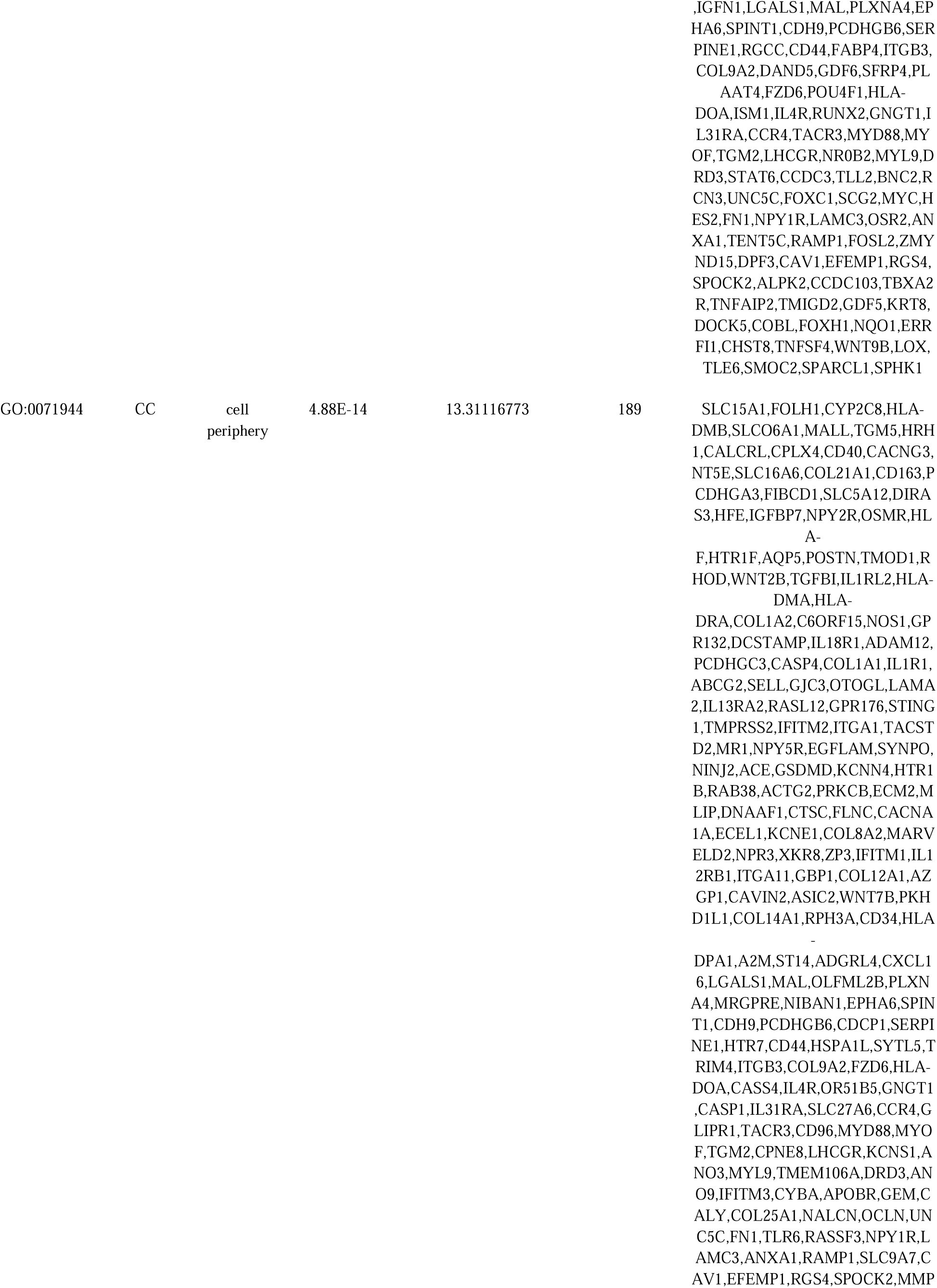
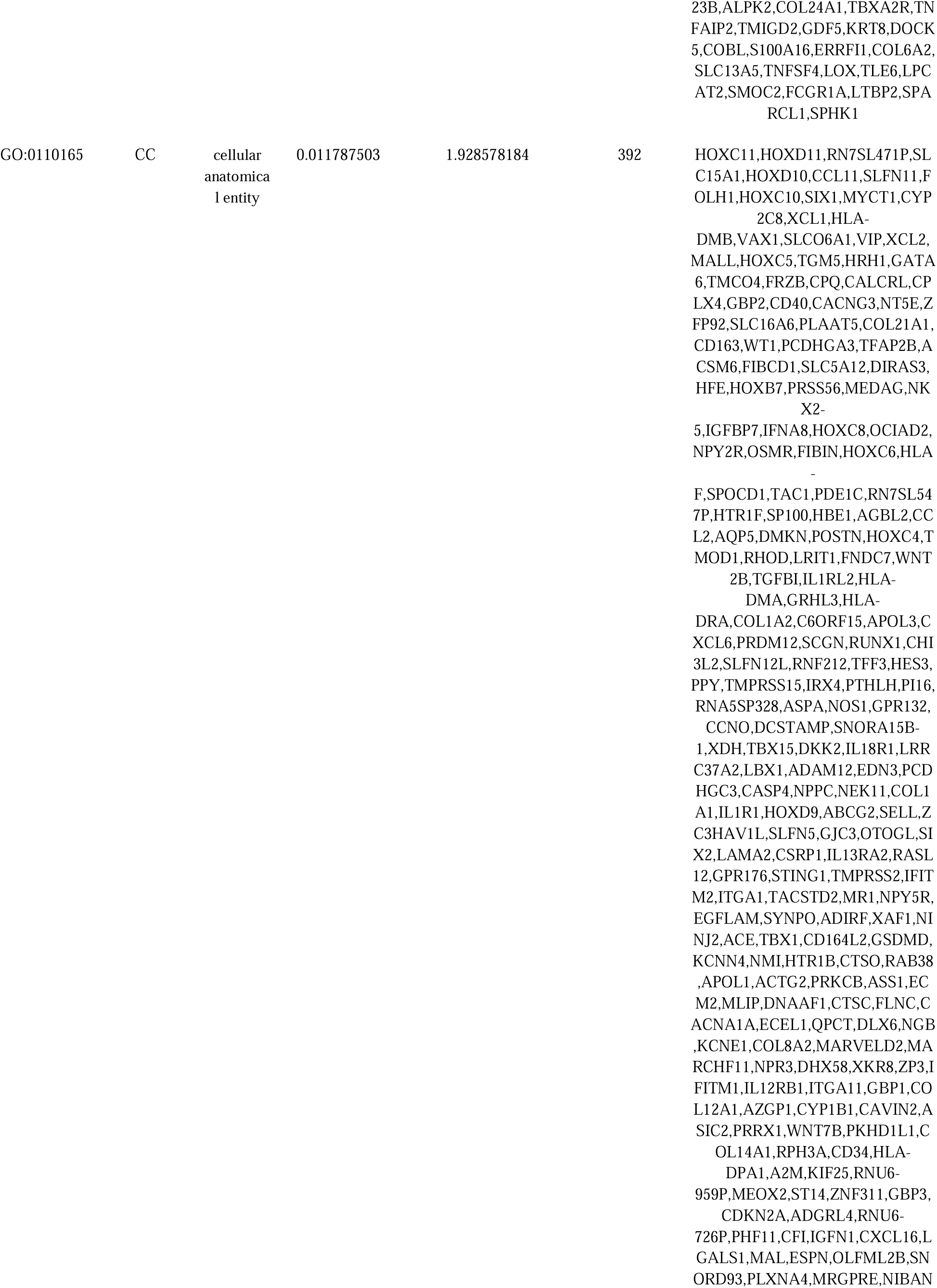
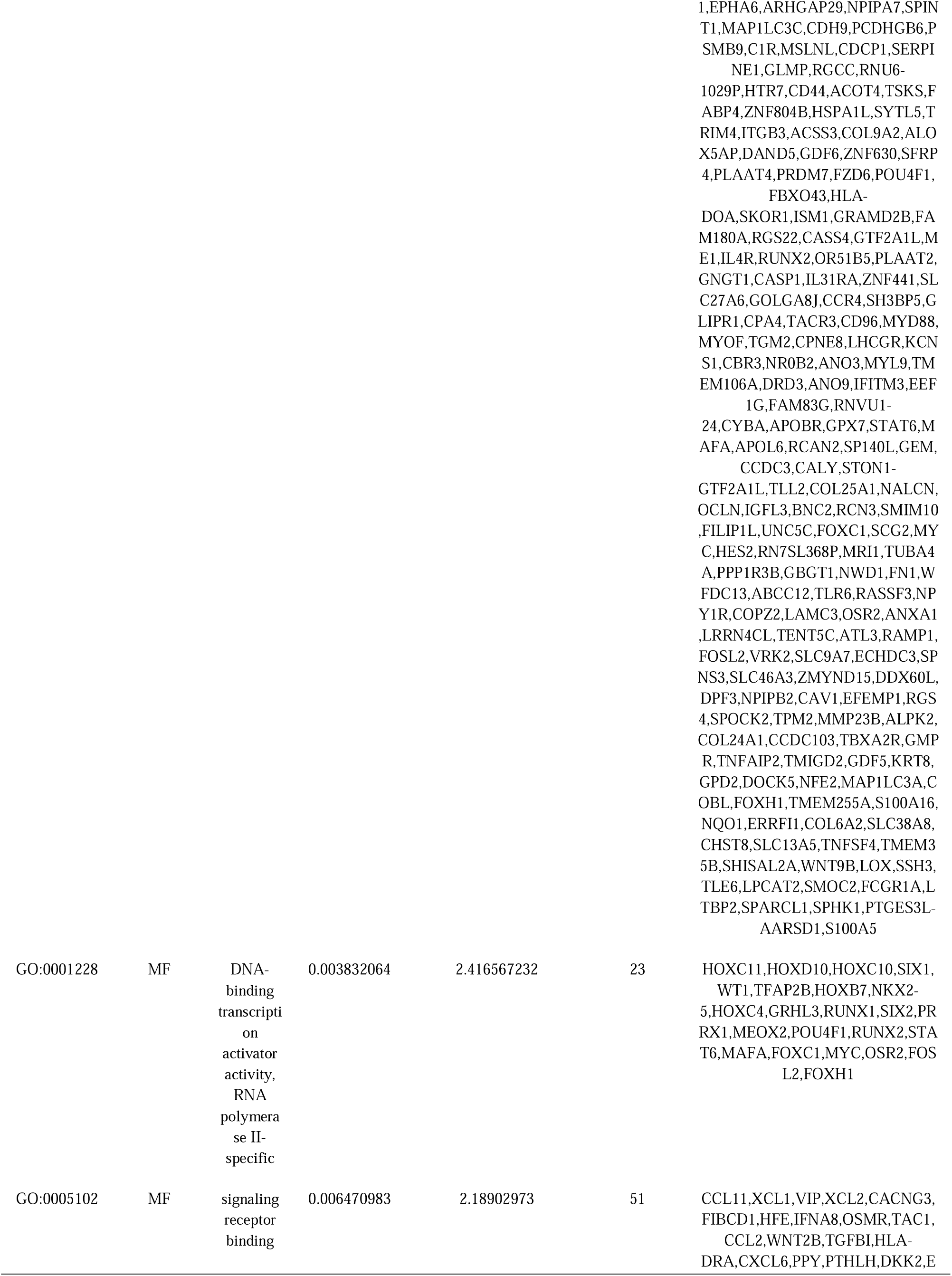
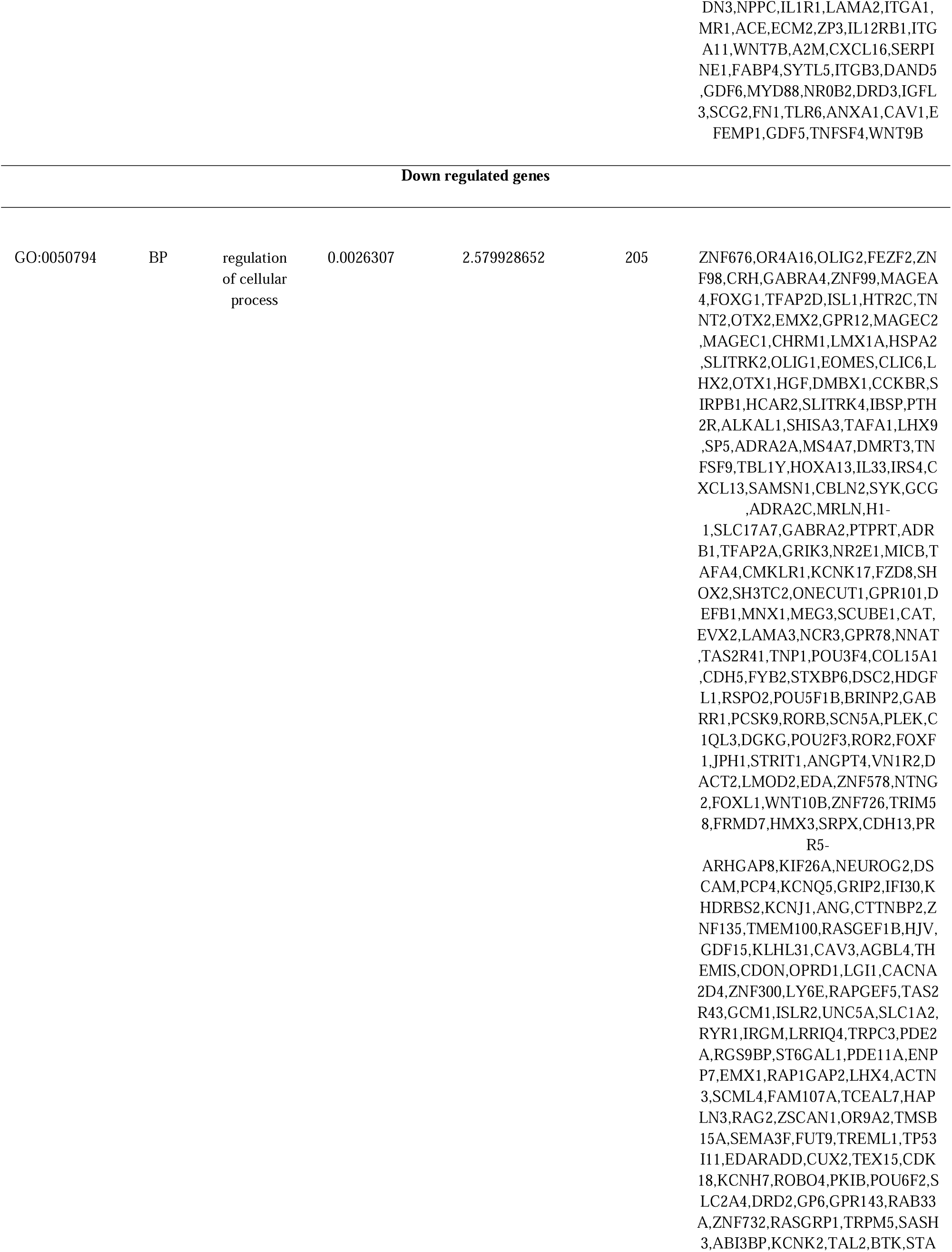
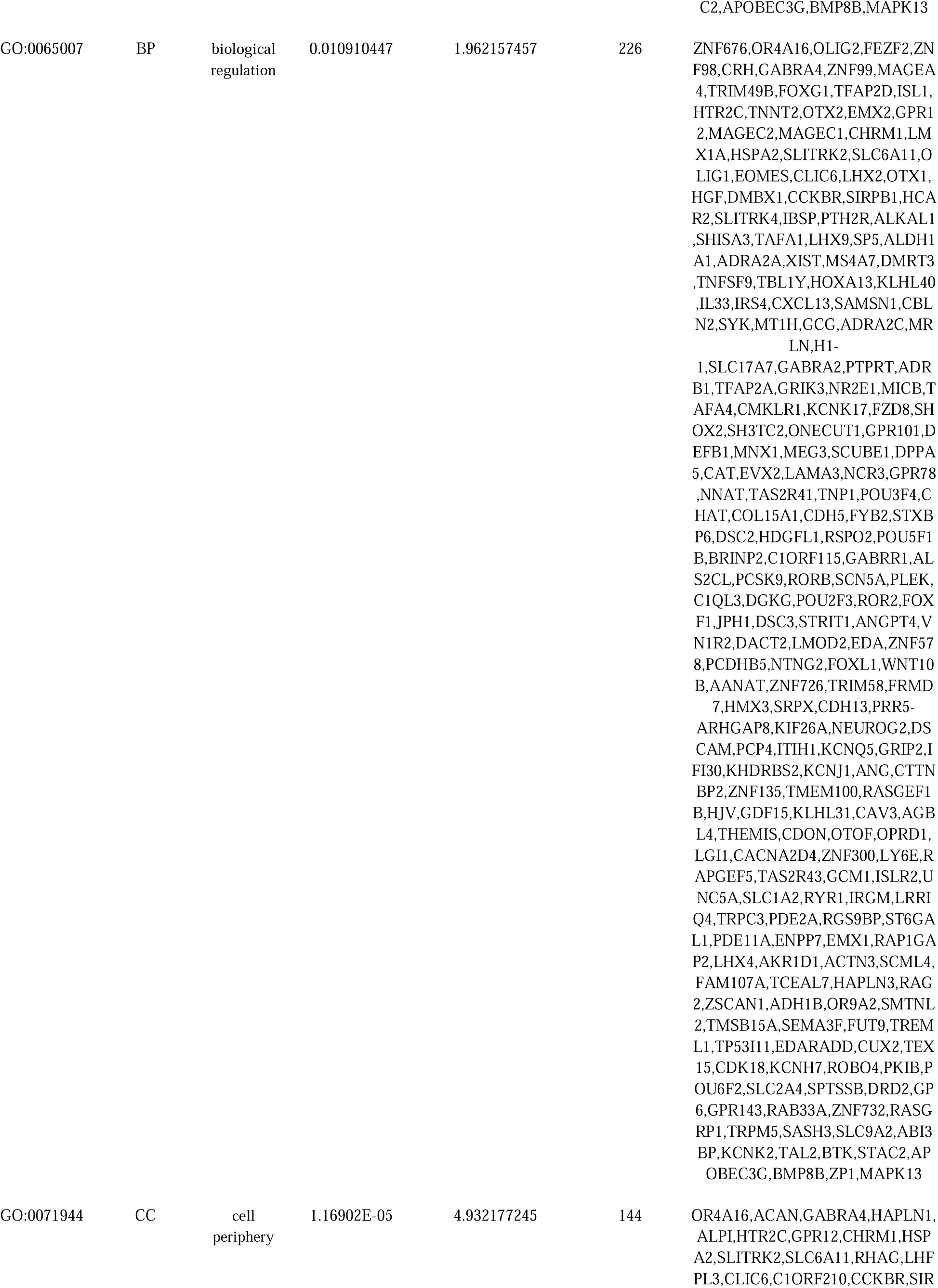
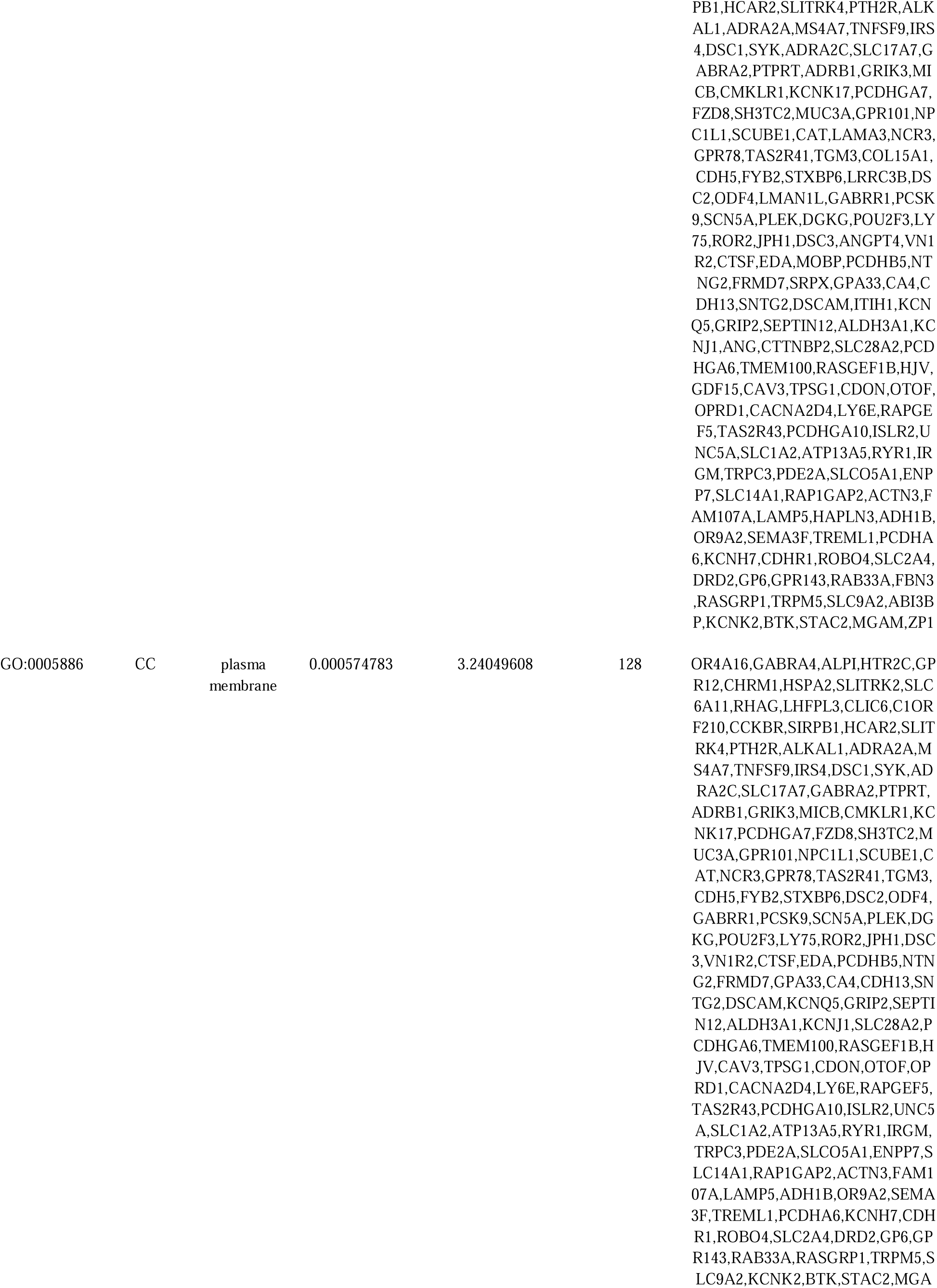
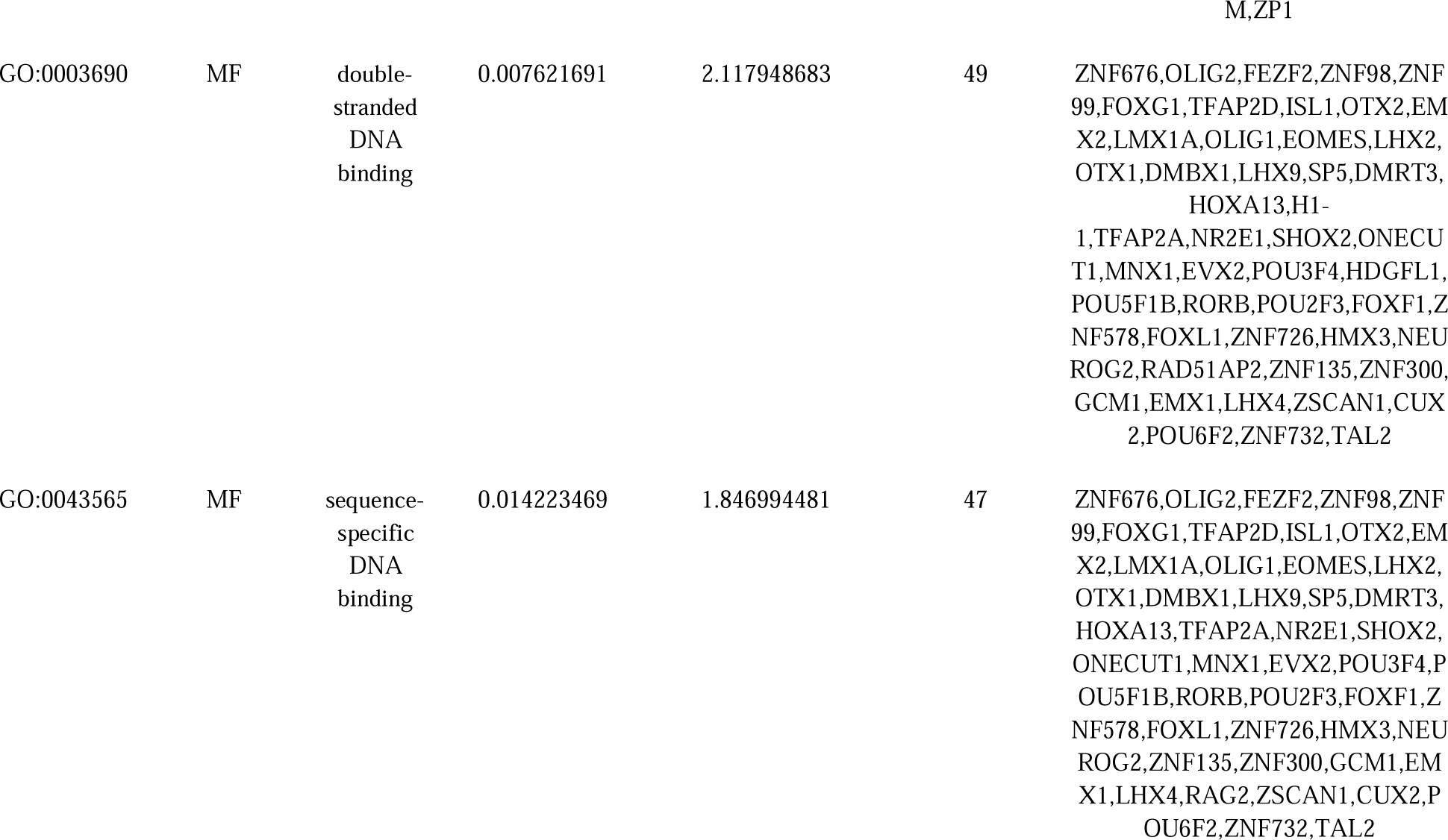
The enriched GO terms of the up and down regulated differentially expressed genes.

**Table 3.** The enriched pathway terms of the up and down regulated differentially expressed genes.

| Pathway ID | Pathway Name | adjusted_p_value | negative_log10_of_adjusted_p_value | Gene Count | Gene |
| --- | --- | --- | --- | --- | --- |
| Up regulated genes |  |  |  |  |  |
| REAC:R-HSA-500792 | GPCR ligand binding | 6.88E-07 | 6.162301143 | 32 | CCL11,XCL1,VIP,XCL2,HRH1,CALCRL,NPY2R,TAC1,HTR1F,CCL2,WNT2B,CXCL6,PPY,PTHLH,GPR132,EDN3,NPY5R,HTR1B,WNT7B,CXCL16,HTR7,FZD6,GNGT1,CCR4,TACR3,LHCGR,DRD3,NPY1R,ANXA1,RAMP1,TBXA2R,WNT9B |
| REAC:R-HSA-1474244 | Extracellular matrix organization | 6.88E-07 | 6.162301143 | 25 | COL21A1,COL1A2,ADAM12,COL1A1,LAMA2,ITGA1,COL8A2,ITGA11,COL12A1,COL14A1,A2M,SERPINE1,CD44,ITGB3,COL9A2,TLL2,COL25A1,FN1,LAMC3,EFEMP1,COL24A1,GDF5,COL6A2,LOX,LTBP2 |
| REAC:R-HSA-373076 | Class A/1 (Rhodopsin-like receptors) | 2.68237E-05 | 4.571481589 | 23 | CCL11,XCL1,XCL2,HRH1,NPY2R,TAC1,HTR1F,CCL2,CXCL6,PPY,GPR132,EDN3,NPY5R,HTR1B,CXCL16,HTR7,CCR4,TACR3,LHCGR,DRD3,NPY1R,A |
|  |  |  |  |  | NXA1,TBXA2R |
| REAC:R-HSA-1280215 | Cytokine Signaling in Immune system | 0.0002394 | 3.620875149 | 34 | CCL11,GBP2,IFNA8,OSMR,HLA-F,SP100,CCL2,IL1RL2,HLA-DRA,COL1A2,IL18R1,IL1R1,IL13RA2,IFITM2,XAF1,GSDMD,IFITM1,IL12RB1,GBP1,HLA-DPA1,GBP3,PSMB9,CD44,IL4R,CASP1,IL31RA,MYD88,IFITM3,STAT6,MYC,FN1,ANXA1,TNFSF4,FCGR1A |
| REAC:R-HSA-375276 | Peptide ligand-binding receptors | 0.000700899 | 3.154344276 | 15 | CCL11,XCL1,XCL2,NPY2R,TAC1,CCL2,CXCL6,PPY,EDN3,NPY5R,CXCL16,CCR4,TACR3,NPY1R,ANXA1 |
| REAC:R-HSA-913531 | Interferon Signaling | 0.002127017 | 2.672228974 | 14 | GBP2,IFNA8,HLA-F,SP100,HLA-DRA,IFITM2,XAF1,IFITM1,GBP1,HLA-DPA1,GBP3,CD44,IFITM3,FCGR1A |
| **Down regulated genes** |  |  |  |  |  |
| REAC:R-HSA-375280 | Amine ligand-binding receptors | 0.001186388 | 2.925773188 | 7 | HTR2C,CHRM1,ADRA2A,ADRA2C,ADRB1,DRD2,GPR143 |
| REAC:R-HSA-372790 | Signaling by GPCR | 0.007766553 | 2.10977171 | 25 | CRH,HTR2C,CHRM1,CCKBR,HCAR2,PTH2R,ADRA2A,CXCL13,GCG,ADRA2C,ADRB1,CMKLR1,FZD8,TAS2R41,DGKG,WNT10B,OPRD1,TAS2R43,TRPC3,PDE2A,PDE11A,DRD2,GPR143,RASGRP1,BTK |
| REAC:R-HSA-112316 | Neuronal System | 0.010991897 | 1.95892736 | 17 | GABRA4,SLITRK2,SLC6A11,SLITRK4,SLC17A7,GABRA2,GRIK3,KCNK17,CHAT,GABRR1,KCNQ5,GRIP2,PPFIBP2,KCNJ1,SLC1A2,KCNH7,KCNK2 |
| REAC:R-HSA-416476 | G alpha (q) signalling events | 0.17634231 | 0.753643474 | 9 | HTR2C,CHRM1,CCKBR,GCG,DGKG,TRPC3,GPR143,RASGRP1,BTK |
| REAC:R-HSA-76002 | Platelet activation, signaling and aggregation | 0.24300464 | 0.614385434 | 9 | HGF,ADRA2A,SYK,ADRA2C,PLEK,DGKG,TRPC3,GP6,RASGRP1 |
| REAC:R-HSA-112315 | Transmission across Chemical Synapses | 0.24300464 | 0.614385434 | 9 | GABRA4,SLC6A11,SLC17A7,GABRA2,GRIK3,CHAT,GABRR1,GRIP2,SLC1A2 |

### Construction of the PPI network and module analysis

To investigate the molecular mechanism of schizophrenia from a systematic perspective, PPI network was constructed to explore the relationship between proteins. PPI network was constructed by IMEx for DEGs. There were 3647 nodes and 6307 edges in the visualization network using the Cyctoscape (Fig. 3). The most high node degree, betweenness, stress and closeness genes were MYC, FN1, CDKN2A, EEF1G, CAV1, ONECUT1, SYK, MAPK13, TFAP2A and BTK, and are listed in Table 4. According the degree of importance, we chose 2 significant modules from the PPI network complex for further analysis using Cytotype MCODE. Functional enrichment analysis showed that module 1 consisted of 54 nodes and 56 edges (Fig.4A), which are mainly associated with cytokine signaling in immune system and multicellular organismal process, and that module 2 consisted of 99 nodes and 117 edges (Fig.4B), which are mainly associated with regulation of cellular process and platelet activation, signaling and aggregation.

**Fig. 3.**
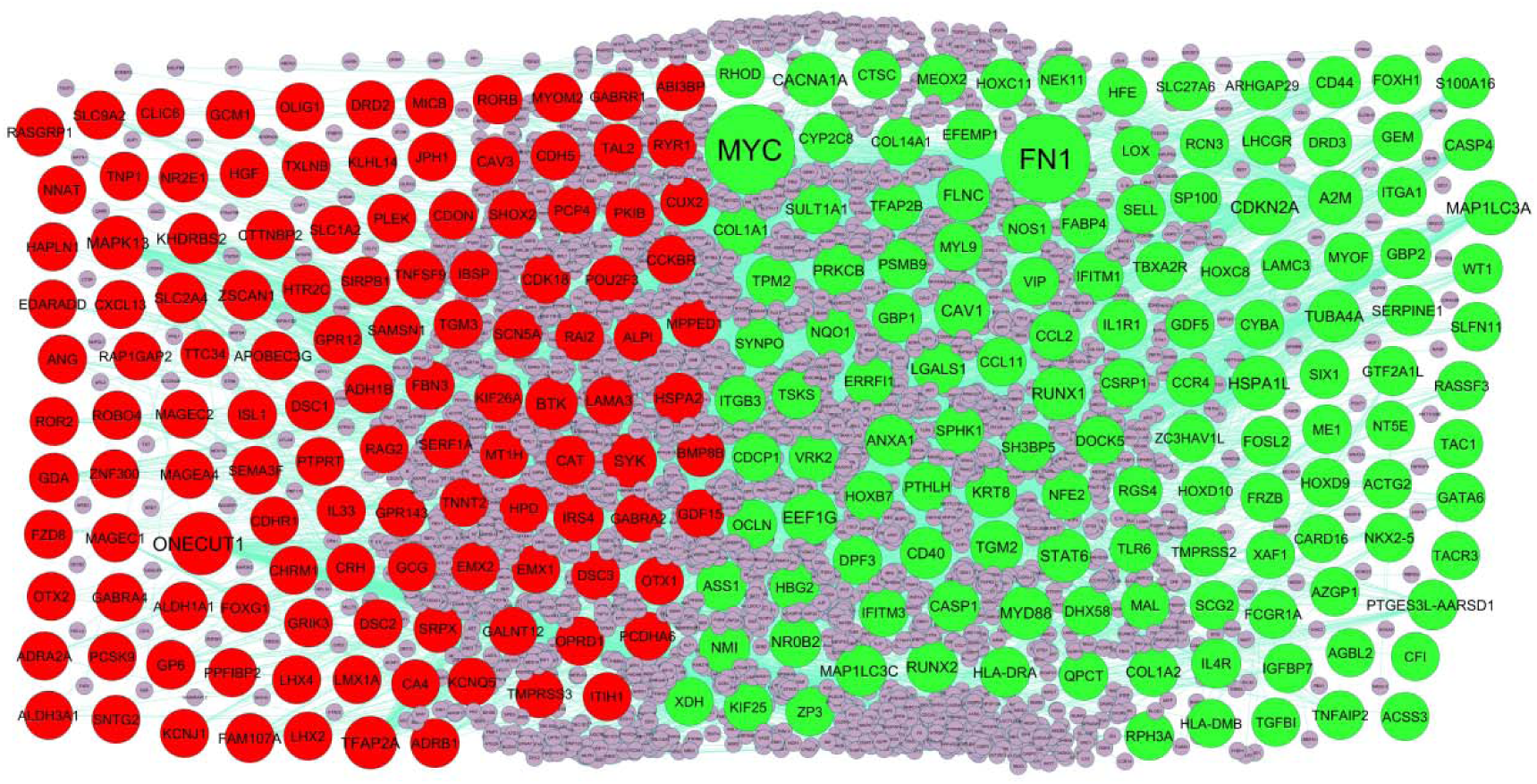
**PPI** network of DEGs. Up regulated genes are marked in parrot green; down regulated genes are marked in red.

**Fig. 4.**
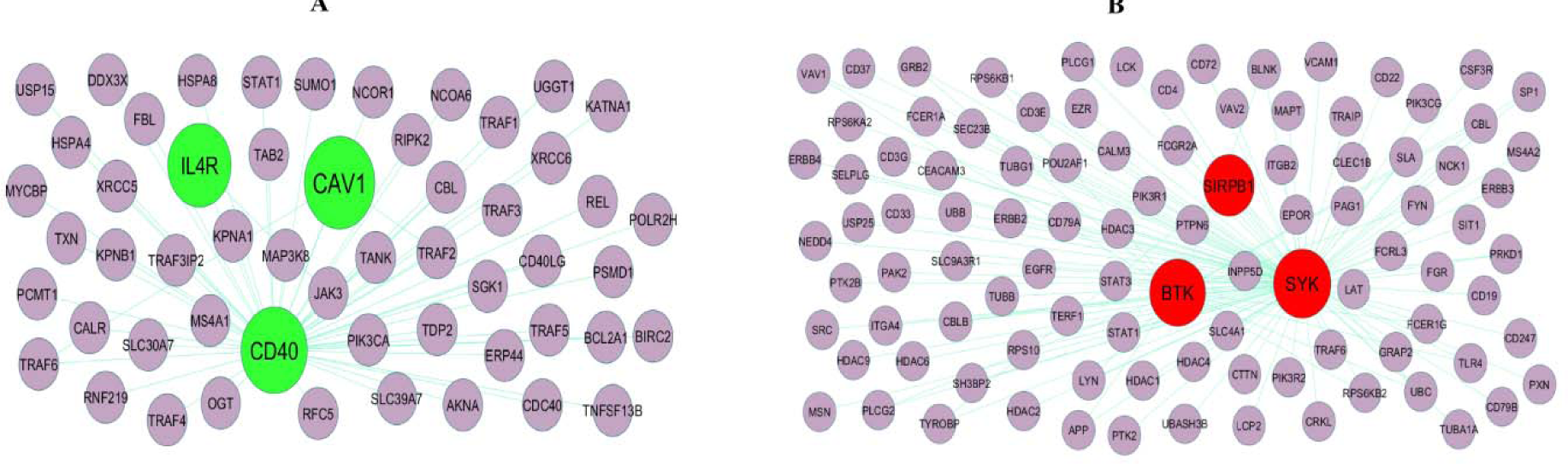
Modules selected from the PPI network. (A) The most significant module was obtained from PPI network with 54 nodes and 56 edges for up regulated genes (B) The most significant module was obtained from PPI network with 99 nodes and 117 edges for down regulated genes. Up regulated genes are marked in parrot green; down regulated genes are marked in red.

**Table 4.**
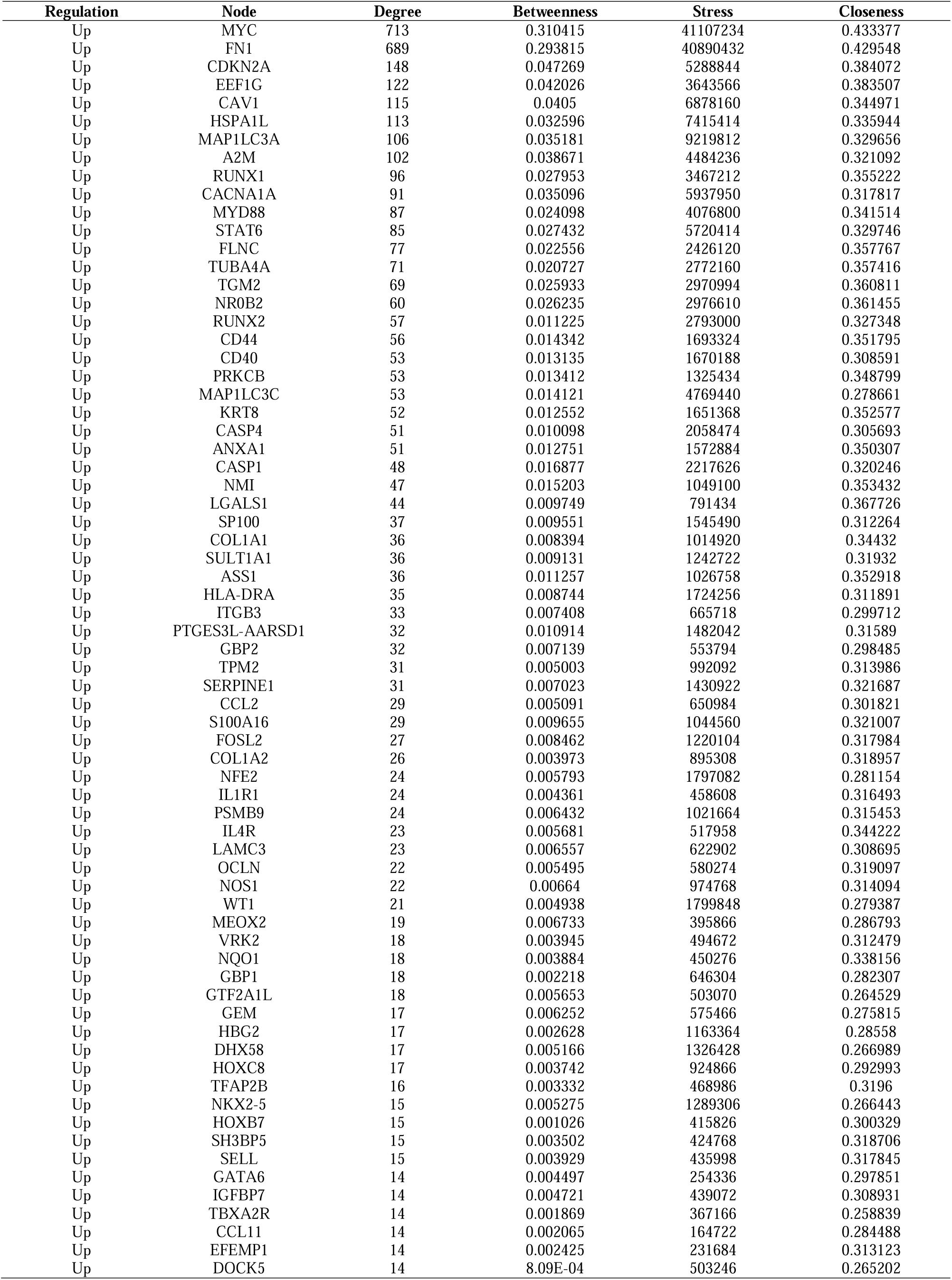
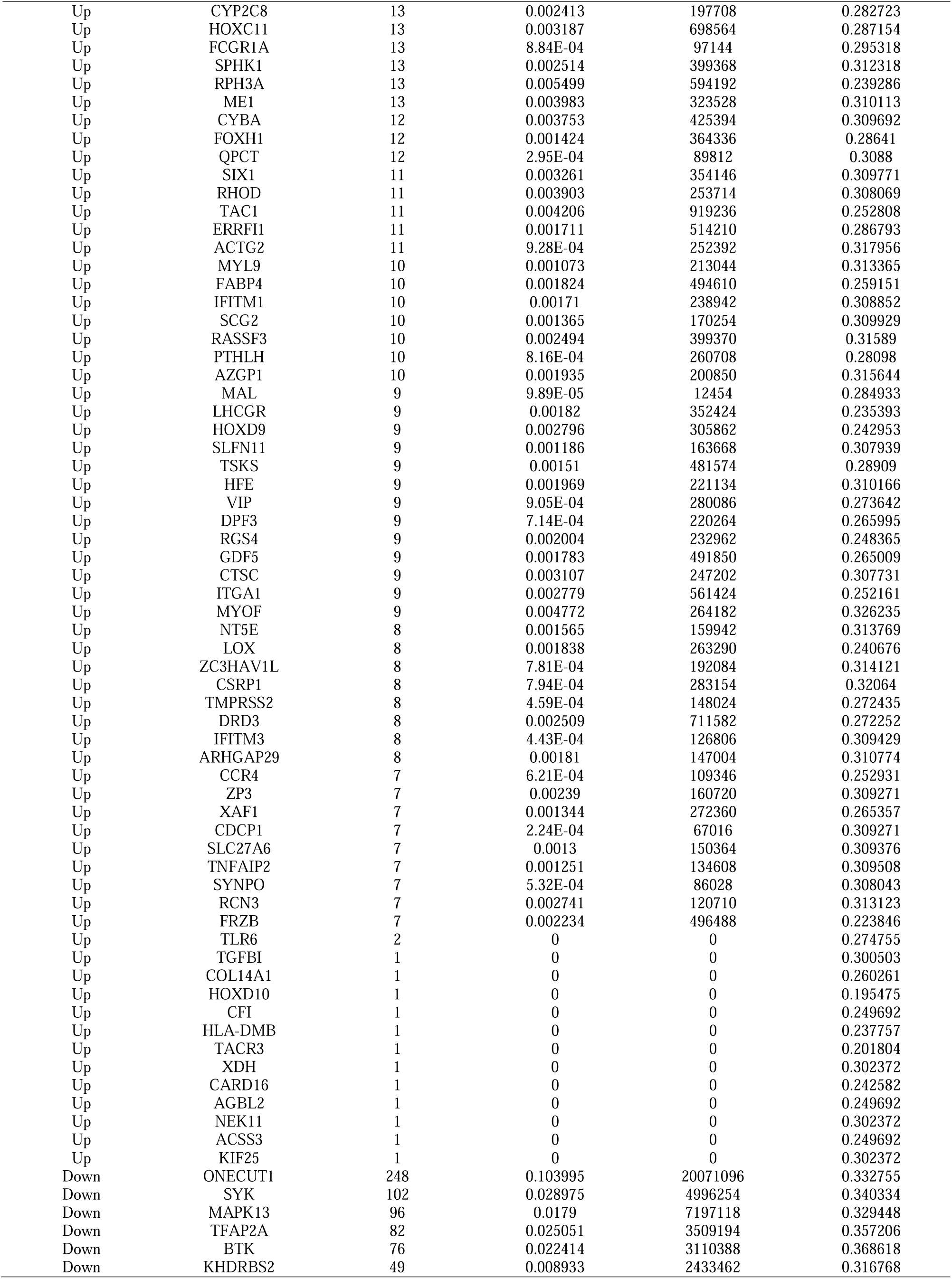
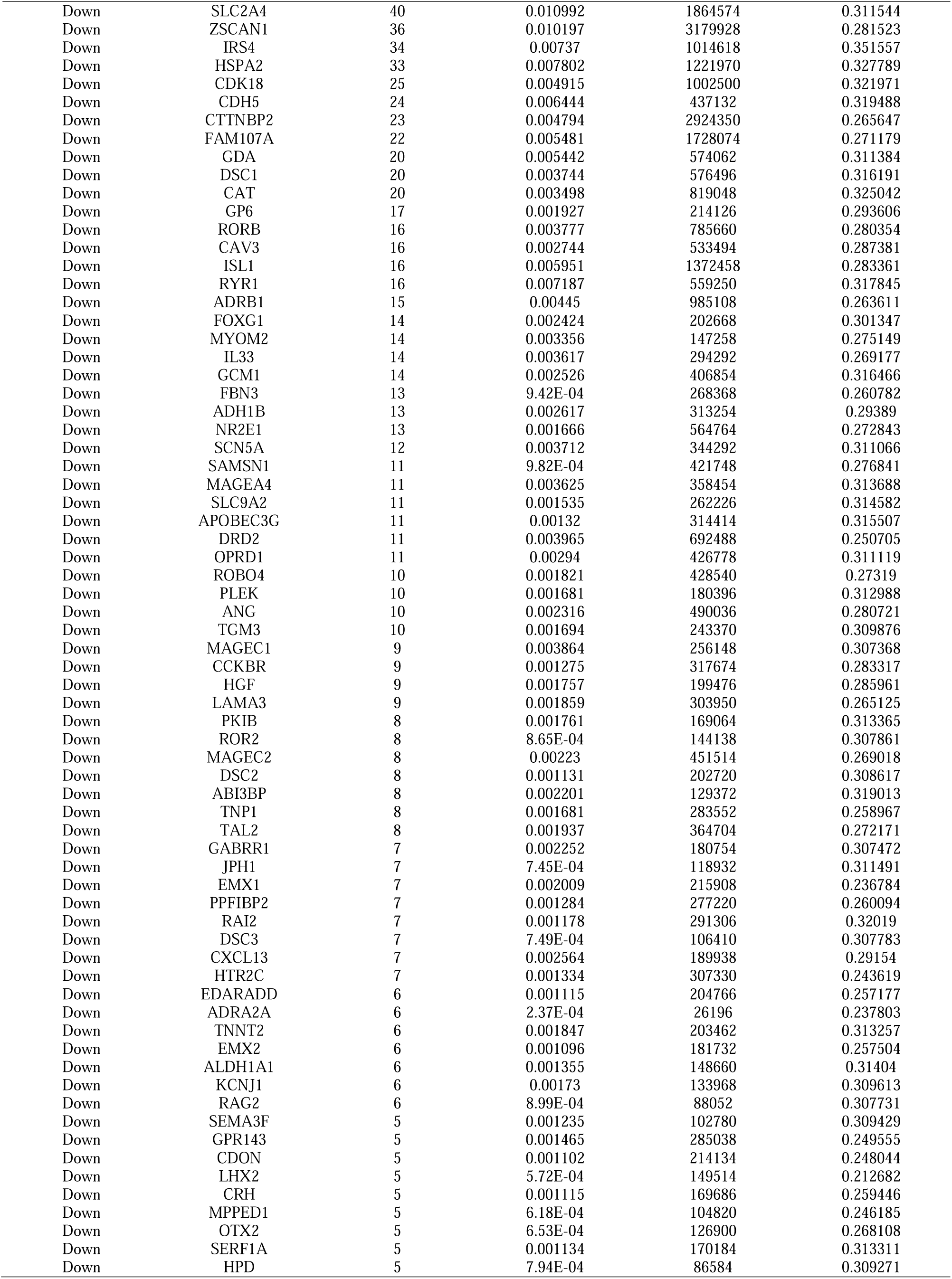
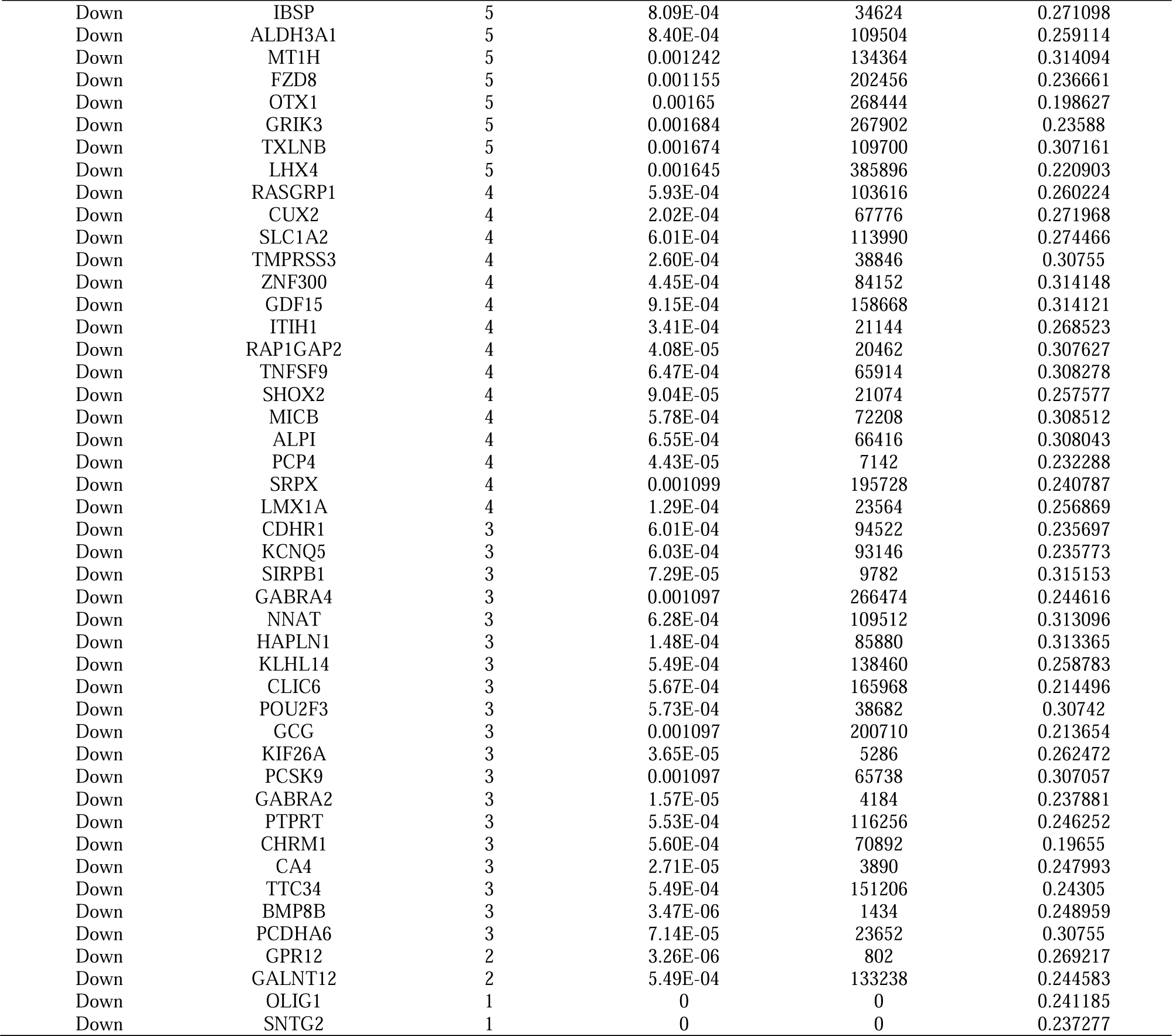
Topology table for up and down regulated genes.

### Construction of the miRNA-hub gene regulatory network

To explore the interactions between schizophrenia related hub genes and miRNA, the miRNA-hub gene regulatory network containing 2191 nodes and 8938 edges was constructed (Fig.5). Of all the nodes, 1937 nodes were miRNAs, while the other 254 nodes were hub genes. The top hub genes for miRNAs were MYC (modulated by 194 miRNAs (ex: hsa-mir-3157-5p)), RUNX1(modulated by 125 miRNAs (ex: hsa-mir-4530)), CAV1 (modulated by 115 miRNAs (ex: hsa-mir-4796-3p)), FN1 (modulated by 105 miRNAs (ex: hsa-mir-132-3p)), CACNA1A (modulated by 69 miRNAs (ex: hsa-mir-10b-5p)), IRS4 (modulated by 134 miRNAs (ex: hsa-mir-769-3p)), TFAP2A (modulated by 100 miRNAs (ex: hsa-mir-4713-5p)), HSPA2 (modulated by 45 miRNAs (ex: hsa-mir-155-5p)), GDA (modulated by 45 miRNAs (ex: hsa-mir-191-5p)) and MAPK13 (modulated by 43 miRNAs (ex: hsa-mir-4516)), and are listed in Table 5.

**Fig. 5.**
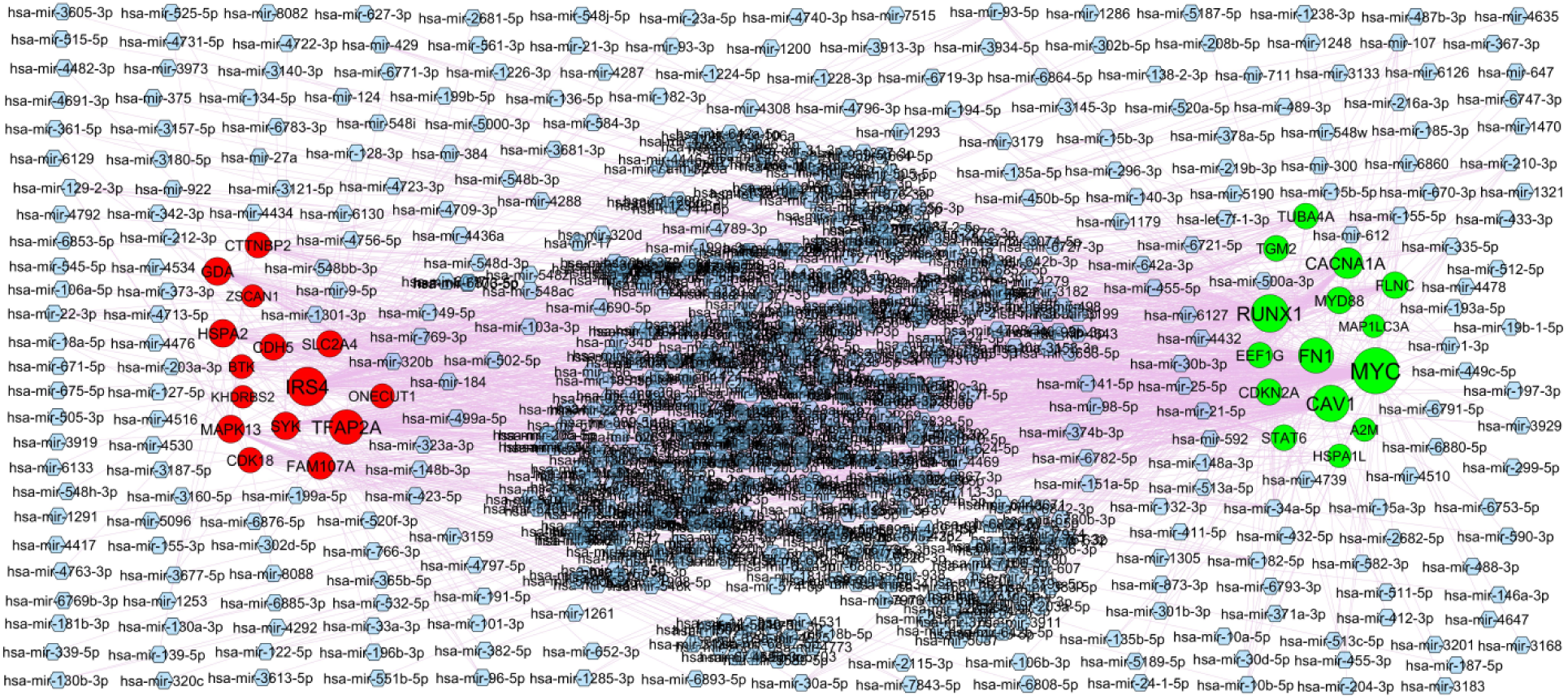
Hub gene - miRNA regulatory network. The blue color diamond nodes represent the key miRNAs; up regulated genes are marked in green; down regulated genes are marked in red.

**Table 5.** MiRNA - hub gene and TF – hub gene topology table.

| Regulation | Hub Genes | Degree | MicroRNA | Regulation | Hub Genes | Degree | TF |
| --- | --- | --- | --- | --- | --- | --- | --- |
| Up | MYC | 194 | hsa-mir-3157-5p | Up | STAT6 | 16 | USF2 |
| Up | RUNX1 | 125 | hsa-mir-4530 | Up | CDKN2A | 16 | SRY |
| Up | CAV1 | 115 | hsa-mir-4796-3p | Up | CAV1 | 16 | HOXA5 |
| Up | FN1 | 105 | hsa-mir-132-3p | Up | FN1 | 14 | RELA |
| Up | CACNA1A | 69 | hsa-mir-10b-5p | Up | MAP1LC3A | 14 | JUND |
| Up | MYD88 | 35 | hsa-mir-384 | Up | RUNX1 | 14 | IRF2 |
| Up | FLNC | 35 | hsa-mir-129-2-3p | Up | FLNC | 12 | PRRX2 |
| Up | CDKN2A | 31 | hsa-mir-191-5p | Up | MYD88 | 9 | SOX10 |
| Up | STAT6 | 28 | hsa-mir-127-5p | Up | CACNA1A | 9 | GATA2 |
| Up | TGM2 | 27 | hsa-mir-107 | Up | A2M | 7 | HNF4A |
| Up | TUBA4A | 27 | hsa-mir-182-5p | Up | MYC | 7 | CEBPB |
| Up | EEF1G | 25 | hsa-mir-342-3p | Up | TUBA4A | 7 | JUN |
| Up | HSPA1L | 10 | hsa-mir-101-3p | Up | HSPA1L | 7 | SREBF1 |
| Up | A2M | 10 | hsa-mir-128-3p | Up | TGM2 | 7 | ELK4 |
| Up | MAP1LC3A | 6 | hsa-mir-22-3p | Up | EEF1G | 2 | ESR1 |
| Down | IRS4 | 134 | hsa-mir-769-3p | Down | GDA | 13 | GATA3 |
| Down | TFAP2A | 100 | hsa-mir-4713-5p | Down | SYK | 13 | FOXC1 |
| Down | HSPA2 | 45 | hsa-mir-155-5p | Down | TFAP2A | 13 | PRDM1 |
| Down | GDA | 45 | hsa-mir-191-5p | Down | IRS4 | 11 | PAX2 |
| Down | MAPK13 | 43 | hsa-mir-4516 | Down | KHDRBS2 | 10 | PDX1 |
| Down | SYK | 42 | hsa-mir-4722-3p | Down | ZSCAN1 | 9 | REL |
| Down | FAM107A | 39 | hsa-mir-146a-3p | Down | MAPK13 | 9 | FOS |
| Down | SLC2A4 | 35 | hsa-mir-1286 | Down | ONECUT1 | 8 | ARID3A |
| Down | CDH5 | 34 | hsa-mir-128-3p | Down | SLC2A4 | 8 | MAX |
| Down | CTTNBP2 | 27 | hsa-mir-15b-5p | Down | CDK18 | 7 | NFIC |
| Down | CDK18 | 24 | hsa-mir-6893-5p | Down | CDH5 | 7 | USF1 |
| Down | ONECUT1 | 16 | hsa-mir-378a-5p | Down | CTTNBP2 | 6 | TEAD1 |
| Down | BTK | 13 | hsa-mir-34a-5p | Down | FAM107A | 4 | NFKB1 |
| Down | KHDRBS2 | 4 | hsa-mir-148b-3p | Down | HSPA2 | 4 | E2F1 |
| Down | ZSCAN1 | 3 | hsa-mir-212-3p | Down | BTK | 2 | SOX5 |

### Construction of the TF-hub gene regulatory network

To explore the interactions between schizophrenia related hub genes and TF, the TF-hub gene regulatory network containing 334 nodes and 1972 edges was constructed (Fig.6). Of all the nodes, 82 nodes were TFs, while the other 252 nodes were hub genes. The top hub genes for TFs were STAT6 (modulated by 16 TFs (ex: USF2)), CDKN2A (modulated by 16 TFs (ex: SRY)), CAV1 (modulated by 16 TFs (ex: HOXA5)), FN1 (modulated by 14 TFs (ex: RELA)), MAP1LC3A (modulated by 14 TFs (ex: JUND)), GDA (modulated by 13 TFs (ex: GATA3)), SYK (modulated by 13 TFs (ex: FOXC1)), TFAP2A (modulated by 13 TFs (ex: PRDM1)), IRS4 (modulated by 11 TFs (ex: PAX2)) and KHDRBS2 modulated by 10 TFs (ex: PDX1)), and are listed in Table 5.

**Fig. 6.**
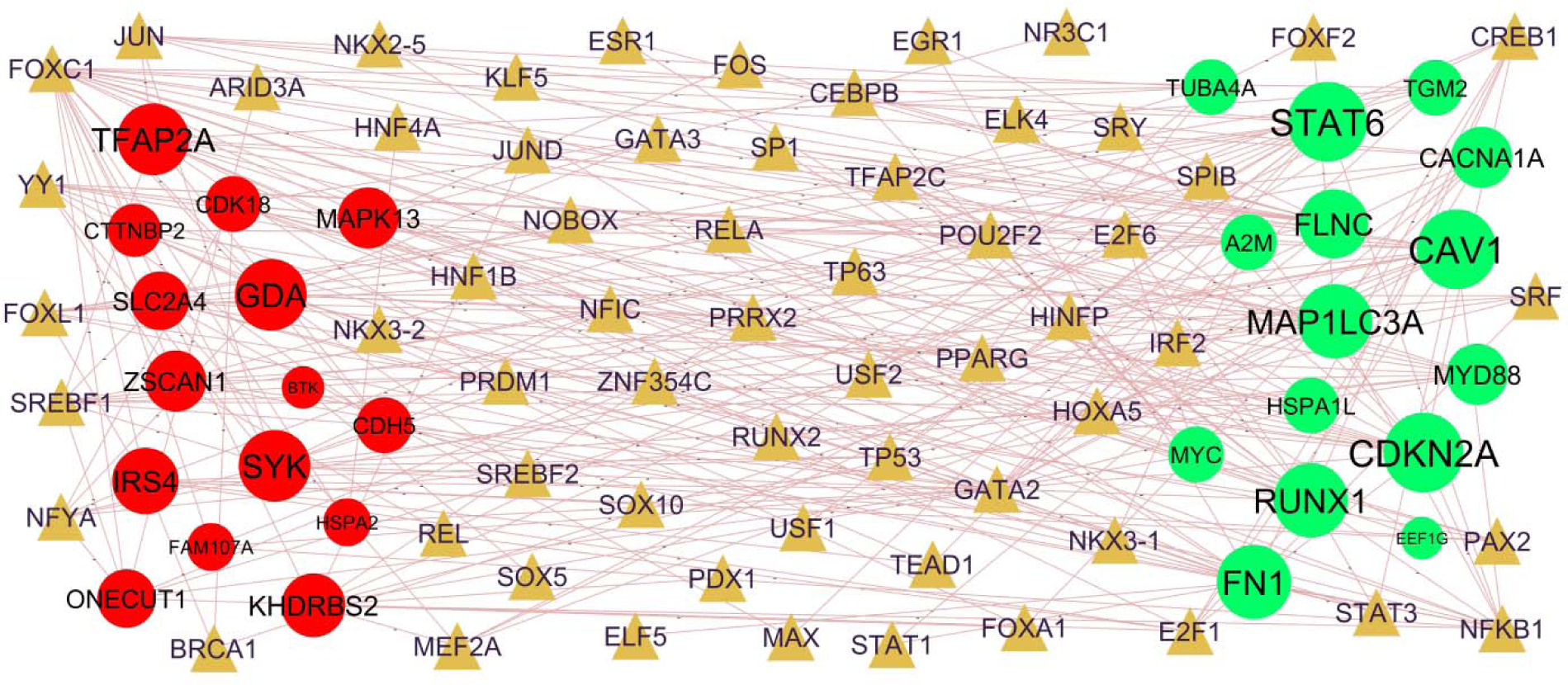
Hub gene - TF regulatory network. The olive color triangle nodes represent the key TFs; up regulated genes are marked in green; down regulated genes are marked in red.

### Receiver operating characteristic curve (ROC) analysis

The AUC values of the ten hub genes were evaluated by ROC curve analysis to examine their sensitivity and specificity for the diagnosis of schizophrenia. All ten hub genes (MYC, FN1, CDKN2A, EEF1G, CAV1, ONECUT1, SYK, MAPK13, TFAP2A and BTK) had AUC values more than 0.8, indicating that they have a strong diagnostic value for schizophrenia (Fig.7).

**Fig. 7.**
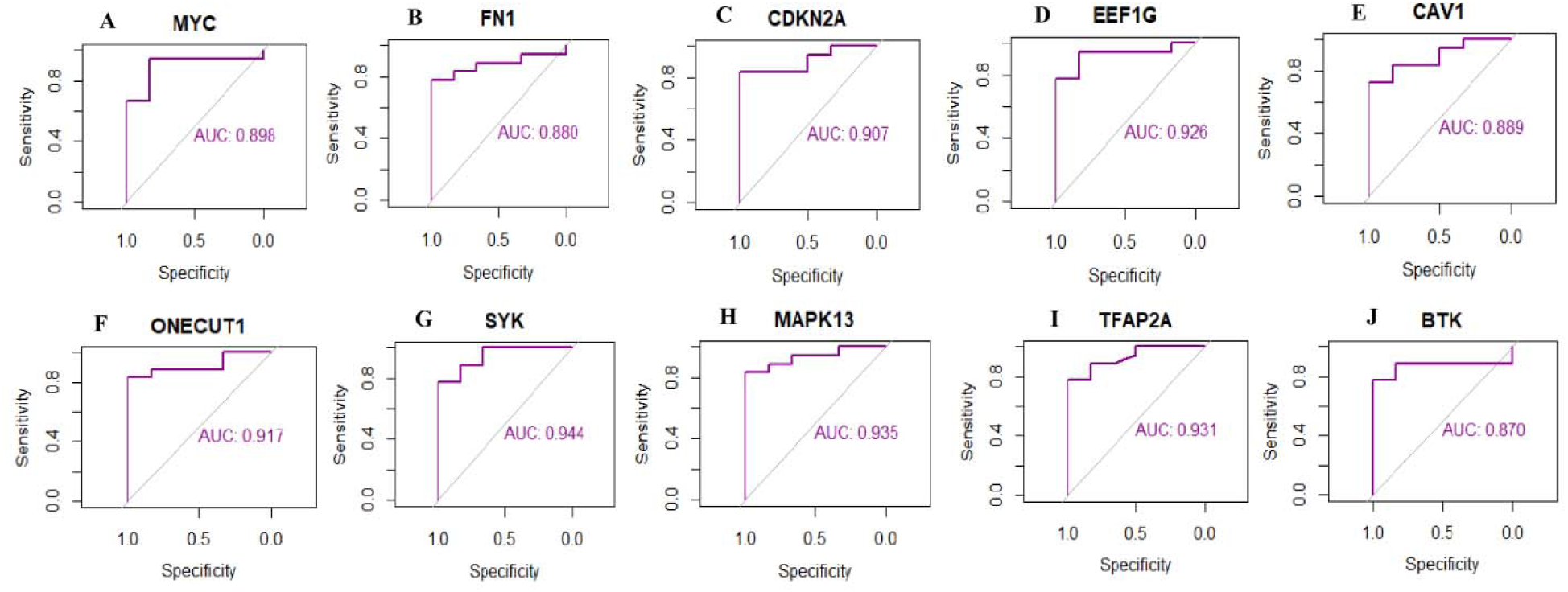
ROC curve analyses of hub genes. A) MYC B) FN1C) CDKN2A D) EEF1G E) CAV1 F) ONECUT1G) SYK H) MAPK13 I) TFAP2A J) BTK

## Discussion

Although numerous relevant investigation of schizophrenia has been performed, early diagnosis, efficacy of treatment and prognosis for schizophrenia remain poorly resolved. For diagnosis and treatment, it is necessary to further understand the molecular pathogenesis resulting in occurrence and advancement. Due to the advancement of NGS technology, the genetic modifications due to disease progression can be detected, indicating gene targets for diagnosis, therapy and prognosis of specific diseases [38-39].

We firstly explored the DEGs in schizophrenia vs. normal control using GSE106589. As a result, a total of 955 DEGs were identified. The expression of HOTAIR (HOX transcript antisense RNA) [40], CCL11 [41], OLIG2 [42] and CRH (corticotropin releasing hormone) [43] might be associated with schizophrenia progression. The abnormal expression of HOTAIR (HOX transcript antisense RNA) [44] might be related to the progression of bipolar disorder. The abnormal expression of HOTAIR (HOX transcript antisense RNA) [45] contributes to the progression of Parkinson’s disease. HOTAIR (HOX transcript antisense RNA) [46] and CCL11 [47] are master regulator that are activated in cardiovascular diseases. HOTAIR (HOX transcript antisense RNA) [48], SLC15A1 [49] and CRH (corticotropin releasing hormone) [50] genes might be related to the pathophysiology of obesity. HOTAIR (HOX transcript antisense RNA) [51], CCL11 [52] and CRH (corticotropin releasing hormone) [53] expression is related to the patients with diabetes mellitus. A studies suggested that HOXD10 [54], CCL11 [55], OLIG2 [56] and CRH (corticotropin releasing hormone) [57] can promote Alzheimer’s disease. CRH (corticotropin releasing hormone) [58] is involved in growth and development of neurodegenerative diseases. Research has shown that CRH (corticotropin releasing hormone) [59] plays an important role in the pathogenesis of hypertension. Result suggests that these significant DEGs play a key role in the progression of schizophrenia.

GO and REACTOME pathway enrichment analyses were used to explore the molecular mechanisms of the enriched genes involved in the occurrence and development of schizophrenia. GPCR ligand binding [60], extracellular matrix organization [61], cytokine signaling in immune system [62], interferon signaling [63], signaling by GPCR [64], neuronal system [65] and platelet activation, signaling and aggregation [66] plays an important role in the schizophrenia. Studies have revealed that SIX1 [67], VIP (vasoactive intestinal peptide) [68], GATA6 [69], FRZB (frizzled related protein) [70], CD40 [71], WT1 [72], PCDHGA3 [73], TFAP2B [74], HFE (homeostatic iron regulator) [75], NKX2-5 [76], IGFBP7 [77], HLA-F [78], CCL2 [79], COL1A2 [80], RUNX1 [81], TFF3 [82], IRX4 [83], NOS1 [84], DKK2 [85], IL18R1 [86], ADAM12 [87], NPPC (natriuretic peptide C) [88], COL1A1 [89], ABCG2 [90], SIX2 [91], CSRP1 [92], MR1 [93], NINJ2 [94], ACE (angiotensin I converting enzyme) [95], TBX1 [96], CTSC (cathepsin C) [97], DLX6 [98], KCNE1 [99], AZGP1 [100], CYP1B1 [101], PRRX1 [102], CD34 [103], A2M [104], CDKN2A [105], SERPINE1 [106], CD44 [107], FABP4 [108], ITGB3 [109], ALOX5AP [110], DAND5 [111], SFRP4 [112], RUNX2 [113], TACR3 [114], MYD88 [115], CYBA (cytochrome b-245 alpha chain) [116], STAT6 [117], FOXC1 [118], FN1 [119], TLR6 [120], CAV1 [121], RGS4 [122], TPM2 [123], TNFSF4 [124], LOX (lysyl oxidase) [125], SMOC2 [126], SPHK1 [127], FOLH1 [128], CYP2C8 [129], CD163 [130], DIRAS3 [131], OSMR (oncostatin M receptor) [132], POSTN (periostin) [133], SELL (selectin L) [134], TMPRSS2 [135], FLNC (filamin C) [136], CXCL16 [137], APOBR (apolipoprotein B receptor) [138], COL6A2 [139], LTBP2 [140], SPARCL1 [141], FOSL2 [142], ISL1 [143], HTR2C [144], TNNT2 [145], HGF (hepatocyte growth factor) [146], IL33 [147], SYK (spleen associated tyrosine kinase) [148], ADRB1 [149], CMKLR1 [150], SHOX2 [151], MEG3 [152], SCUBE1 [153], CAT (catalase) [154], LAMA3 [155], COL15A1 [156], DSC2 [157], RSPO2 [158], PCSK9 [159], SCN5A [160], FOXF1 [161], DACT2 [162], LMOD2 [163], CDH13 [164], DSCAM (DS cell adhesion molecule) [165], PCP4 [166], ANG (angiogenin) [167], GDF15 [168]. RYR1 [169], IRGM (immunity related GTPase M) [170], TRPC3 [171], PDE2A [172], SCML4 [173], SEMA3F [155], CUX2 [174], ROBO4 [175], DRD2 [176], GP6 [177], TRPM5 [178], ABI3BP [179], ACAN (aggrecan) [180] and NPC1L1 [181] plays a key role in cardiovascular diseases. Previous studies have reported that the XCL1 [182], HLA-DMB [183], CD40 [184], HLA-DRA [185], RUNX1 [186], IL18R1 [187], NINJ2 [188], ACE (angiotensin I converting enzyme) [189], CD44 [190], IL4R [191], MYD88 [192], WNT9B [193], CXCL16 [194], CXCL13 [195], RORB (RAR related orphan receptor B) [196], GDF15 [197], THEMIS (thymocyte selection associated) [198], KCNH7 [199], BTK (Bruton tyrosine kinase) [200] and MOBP (myelin associated oligodendrocyte basic protein) [201] are a key regulators of multiple sclerosis. Recently, increasing evidence demonstrated that HLA-DMB [202], VIP (vasoactive intestinal peptide) [203], GATA6 [204], CD40 [205], TFAP2B [206], HFE (homeostatic iron regulator) [207], IGFBP7 [208], NPY2R [209], CCL2 [210], AQP5 [211], HLA-DMA [212], RUNX1 [81], PPY (pancreatic polypeptide) [213], ASPA (aspartoacylase) [214], NOS1 [215], ADAM12 [216], NPPC (natriuretic peptide C) [217], COL1A1 [218], IL1R1 [219], ABCG2 [220], ACE (angiotensin I converting enzyme) [221], CD34 [222], HLA-DPA1 [223], A2M [224], MEOX2 [225], CDKN2A [226], SERPINE1 [227], CD44 [228], FABP4 [108], ITGB3 [229], ALOX5AP [230], SFRP4 [231], ISM1 [232], IL4R [233], RUNX2 [234], CASP1 [235], CCR4 [236], MYD88 [237], DRD3 [238], STAT6 [239], ANXA1 [240], CAV1 [241], RGS4 [242], SPHK1 [243], CYP2C8 [244], CD163 [245], DIRAS3 [131], POSTN (periostin) [246], SELL (selectin L) [247], TMPRSS2 [248], CXCL16 [249], FOSL2 [250], ISL1 [251], HGF (hepatocyte growth factor) [252], ADRA2A [253], IL33 [254], SYK (spleen associated tyrosine kinase) [148], GCG (glucagon) [255], PTPRT (protein tyrosine phosphatase receptor type T) [256], GRIK3 [257], NR2E1 [258], CMKLR1 [259], ONECUT1 [260], DEFB1 [261], MNX1 [262], MEG3 [263], CAT (catalase) [264], PCSK9 [265], PLEK (pleckstrin) [266], EDA (ectodysplasin A) [267], KCNJ1 [268], ANG (angiogenin) [269], GDF15 [270], TRPC3 [271], RAG2 [272], ROBO4 [273], SLC2A4 [274], DRD2 [275], GP6 [276], RASGRP1 [277], TRPM5 [278], NPC1L1 [279], ALDH3A1 [280] and ADH1B [281] were altered expressed in diabetes mellitus. Studies had shown that VIP (vasoactive intestinal peptide) [282], HFE (homeostatic iron regulator) [283], IGFBP7 [284], HOXC8 [285], CCL2 [286], GRHL3 [287], NOS1 [288], ADAM12 [289], NINJ2 [290], ACE (angiotensin I converting enzyme) [291], TBX1 [292], HTR1B [293], CD34 [294], HLA-DPA1 [295], HTR7 [296], FABP4 [297], ITGB3 [298], TACR3 [299], MYD88 [300], TGM2 [301], DRD3 [302], CAV1 [303], RGS4 [304], SLCO6A1 [305], CD163 [306], CACNA1A [307], HSPA1L [308], NALCN (sodium leak channel, non-selective) [309], HTR2C [310], CHRM1 [311], LMX1A [312], SLITRK2 [313], CCKBR (cholecystokinin B receptor) [314], ADRA2A [315], IL33 [316], IRS4 [317], ADRA2C [315], GRIK3 [318], NR2E1 [319], MICB (MHC class I polypeptide-related sequence B) [320], MEG3 [321], CAT (catalase) [322], GPR78 [323], PCSK9 [324], SCN5A [325], NTNG2 [326], CDH13 [327], LGI1 [328], SLC1A2 [329], PDE2A [330], KCNH7 [331], DRD2 [332], GPR143 [333], RASGRP1 [334] and ACAN (aggrecan) [335] were associated with schizophrenia. VIP (vasoactive intestinal peptide) [336], CD40 [337], CCL2 [338], DKK2 [339], ACE (angiotensin I converting enzyme) [340], KCNN4 [341], A2M [342], CDKN2A [343], MYD88 [344], TLR6 [345], ANXA1 [346], SPHK1 [347], CACNA1A [348], SLITRK2 [349], IL33 [350], CAT (catalase) [351], GPR78 [352], ANG (angiogenin) [353], GDF15 [354] and ADH1B [355] might be a potential therapeutic targets for neurodegenerative diseases treatment. At present, abnormal expression of VIP (vasoactive intestinal peptide) [356], HFE (homeostatic iron regulator) [357], CCL2 [358], HLA-DRA [359], TFF3 [360], NOS1 [361], NPPC (natriuretic peptide C) [362], ACE (angiotensin I converting enzyme) [363], GSDMD (gasdermin D) [364], A2M [365], PLXNA4 [366], CD44 [367], CASP1 [364], DRD3 [368], UNC5C [369], CAV1 [370], SPHK1 [371], CD163 [372], RPH3A [373], HGF (hepatocyte growth factor) [374], CCKBR (cholecystokinin B receptor) [375], TNFSF9 [376], MEG3 [377], GPR78 [378], NEUROG2 [379], ANG (angiogenin) [380], GDF15 [381], UNC5A [382], SLC1A2 [383], DRD2 [384], GPR143 [385], RASGRP1 [386] and MOBP (myelin associated oligodendrocyte basic protein) [387] have been found in a Parkinson’s disease. VIP (vasoactive intestinal peptide) [388], CD40 [389], WT1 [390], HFE (homeostatic iron regulator) [391], TAC1 [392], AQP5 [393], WNT2B [394], RUNX1 [395], NOS1 [396], DKK2 [339], ADAM12 [397], ABCG2 [398], NINJ2 [399], ACE (angiotensin I converting enzyme) [400], PRKCB (protein kinase C beta) [401], A2M [365], MEOX2 [402], CDKN2A [403], PLXNA4 [404], SPINT1 [405], SERPINE1 [406], RGCC (regulator of cell cycle) [407], CD44 [408], CASP1 [409], MYD88 [410], DRD3 [411], UNC5C [412], LOX (lysyl oxidase) [413], SPHK1 [414], RPH3A [415], CXCL16 [416], CASS4 [417], IFITM3 [418], COL25A1 [419], SPARCL1 [420], FOXG1 [421], CHRM1 [422], HSPA2 [423], HGF (hepatocyte growth factor) [424], IL33 [425], MEG3 [426], RSPO2 [427], PCSK9 [428], PCSK9 [429], RORB (RAR related orphan receptor B) [430], ANGPT4 [431], CDH13 [432], PCP4 [433], ANG (angiogenin) [434], GDF15 [435], OPRD1 [436], PDE11A [437], TREML1 [438], GP6 [439], BTK (Bruton tyrosine kinase) [440], DSC1 [441], LAMP5 [442] and ADH1B [443] were identified to be associated with Alzheimer’s disease. VIP (vasoactive intestinal peptide) [444], CD40 [445], TFAP2B [446], NPY2R [447], CCL2 [448], COL1A2 [80], RUNX1 [449], PPY (pancreatic polypeptide) [213], ASPA (aspartoacylase) [214], TBX15 [450], ADAM12 [451], NPPC (natriuretic peptide C) [88], COL1A1 [89], ABCG2 [452], STING1 [453], NPY5R [454], ACE (angiotensin I converting enzyme) [455], HTR1B [456], PRKCB (protein kinase C beta) [457], NPR3 [458], CYP1B1 [459], CD34 [460], A2M [461], CDKN2A [226], SERPINE1 [227], CD44 [462], FABP4 [463], ALOX5AP [464], RUNX2 [465], CASP1 [466], MYD88 [467], STAT6 [468], TLR6 [469], NPY1R [470], ANXA1 [471], CAV1 [472], RGS4 [473], DOCK5 [474], COBL (cordon-bleu WH2 repeat protein) [475], LOX (lysyl oxidase) [476], SPHK1 [477], CD163 [478], POSTN (periostin) [479], TMPRSS2 [480], CXCL16 [481], IFITM3 [482], ISL1 [483], HTR2C [484], HSPA2 [485], HGF (hepatocyte growth factor) [486], ADRA2A [487], IL33 [486], GCG (glucagon) [255], PTPRT (protein tyrosine phosphatase receptor type T) [256], ADRB1 [489], NR2E1 [490], CMKLR1 [491], MEG3 [492], SCUBE1 [493], CAT (catalase) [264], PCSK9 [159], SCN5A [494], EDA (ectodysplasin A) [267], WNT10B [495], CDH13 [496], GDF15 [168], ACTN3 [497], SLC2A4 [274], DRD2 [498], TRPM5 [278], BMP8B [499], ACAN (aggrecan) [500], NPC1L1 [501] and ADH1B [502] have been shown to be a biomarkers of obesity. VIP (vasoactive intestinal peptide) [503], GATA6 [504], CD40 [505], WT1 [506], HFE (homeostatic iron regulator) [507], IGFBP7 [508], CCL2 [509], AQP5 [510], RUNX1 [511], NOS1 [512], XDH (xanthine dehydrogenase) [513], ADAM12 [87], NPPC (natriuretic peptide C) [514], IL1R1 [515], SIX2 [516], ACE (angiotensin I converting enzyme) [517], CYP1B1 [518], CD34 [519], CDKN2A [105], EPHA6 [520], CD44 [521], FABP4 [522], ITGB3 [523], RUNX2 [524], CASP1 [525], MYD88 [526], DRD3 [527], STAT6 [528], RCN3 [529], FOXC1 [530], TLR6 [531], ANXA1 [532], RAMP1 [533], CAV1 [534], LOX (lysyl oxidase) [535], SPHK1 [536], CYP2C8 [537], CD163 [538], POSTN (periostin) [539], TMPRSS2 [540], RAB38 [541], ECM2 [542], CACNA1A [543], CAVIN2 [544], CXCL16 [545], COL6A2 [546], SPARCL1 [547], HGF (hepatocyte growth factor) [548], CCKBR (cholecystokinin B receptor) [549], ADRA2A [253], IL33 [147], CXCL13 [550], CBLN2 [551], ADRB1 [149], CMKLR1 [552], SCUBE1 [553], CAT (catalase) [554], PCSK9 [555], FOXF1 [556], CDH13 [557], GDF15 [558], TRPC3 [559], GPR143 [560], BTK (Bruton tyrosine kinase) [561], ACAN (aggrecan) [562] and ADH1B [281] have been found to be altered expression in hypertension. FRZB (frizzled related protein) [563], HLA-DRA [564], CD44 [565], CASP1 [566], LOX (lysyl oxidase) [567], ANG (angiogenin) [568] and RAG2 [569] genes expression were found to be elevated in amyotrophic lateral sclerosis. IGFBP7 [570], TFF3 [360], ACE (angiotensin I converting enzyme) [517], ANXA1 [346], EFEMP1 [571], ANGPT4 [431], GDF15 [381] and MOBP (myelin associated oligodendrocyte basic protein) [572] expression are altered in the patients with dementia. NPY2R [573], CASP1 [574], CHRM1 [575] and MEG3 [576] have been proposed as novel biomarkers for Huntington disease. Some studies have shown that SPOCD1 [577], DRD3 [578], DOCK5 [579], SLCO6A1 [305], HTR2C [580], OTX2 [581], LMX1A [582], NR2E1 [319], GPR78 [323], RORB (RAR related orphan receptor B) [583], NTNG2 [326], PRR5-ARHGAP8 [584], CACNA2D4 [585], SLC1A2 [586], CUX2 [587], KCNH7 [588] and ACAN (aggrecan) [335] plays a certain role in bipolar disorder. The altered expression of ABCG2 [589], LAMC3 [590], CACNA1A [591], COL6A2 [592], SLC13A5 [593], GABRA2 [594], SCN5A [160], KCNQ5 [595], SLC1A2 [596], TRPC3 [597], GABRA4 [598], SLC6A11 [599], KCNQ5 [600] and SLCO5A1 [601] are associated with epilepsy. In summary, DEGs involved in GO term and REACTOME pathway were more likely related to schizophrenia and might be important targets in schizophrenia therapy.

To explore the molecular pathogenesis of schizophrenia, we constructed PPI network and module analysis for systematic analysis. In order to further analyze the whole PPI network, the topological analysis was used to explain the importance of the hub genes in the network and the influence of the hub genes on the network. Previous studies had shown that the altered expression of FN1 [119], CDKN2A [105], CAV1 [121], SYK (spleen associated tyrosine kinase) [148], and CD40 [71] were closely related to the occurrence of cardiovascular diseases. A study had shown that regulation of CDKN2A [226], CAV1 [241], ONECUT1 [260], SYK (spleen associated tyrosine kinase) [148], IL4R [233] and CD40 [205] promoted the diabetes mellitus. CDKN2A [343] and CD40 [337] are involved in mediating the progression of neurodegenerative diseases. Previous studies report the altered expression of CDKN2A [403], BTK (Bruton tyrosine kinase) [440] and CD40 [389] in the nervious tissue obtained from Alzheimer’s disease patients. CDKN2A [226], CAV1 [472] and CD40 [445] are involved in the mediation of obesity. The results showed that CDKN2A [105], CAV1 [534], BTK (Bruton tyrosine kinase) [561] and CD40 [505] were expressed in hypertension. CAV1 [303] and IRS4 [317] expression is significantly regulated in schizophrenia patients. Previous studies report that CAV1 [370] is involved in Parkinson’s disease. BTK (Bruton tyrosine kinase) [200], IL4R [191] and CD40 [184] expression is associated with clinical and biochemical markers for multiple sclerosis. We identified MYC (MYC proto-oncogene, bHLH transcription factor), EEF1G, MAPK13, TFAP2A and SIRPB1 might serve as novel biomarkers for schizophrenia. The results suggested that these hub genes might play significant roles in schizophrenia. These findings indicate that hub genes might plays a key role in the molecular pathogenesis of schizophrenia., and are a key biomarkers linking schizophrenia.

To explore the molecular mechanism of schizophrenia, we constructed the schizophrenia related miRNA-hub gene regulatory network and TF-hub gene regulatory network. Moreover, we performed topological analysis and acquired hub genes, miRNAs and TFs with high topological features. RUNX1 [81], CAV1 [121], FN1 [119], STAT6 [117], CDKN2A [105], SYK (spleen associated tyrosine kinase) [148], hsa-mir-769-3p [602], HOXA5 [603], JUND (JunD proto-oncogene, AP-1 transcription factor subunit) [604] and FOXC1 [605] were associated with cardiovascular diseases. RUNX1 [186], hsa-mir-132-3p [606] and HOXA5 [607] was an important therapeutic targets of multiple sclerosis. RUNX1 [81], CAV1 [241], STAT6 [239], CDKN2A [226], SYK (spleen associated tyrosine kinase) [148], hsa-mir-132-3p [608], hsa-mir-10b-5p [609], hsa-mir-155-5p [610], hsa-mir-191-5p [611], SRY (sex determining region Y) [612], PAX2 [613] and PDX1 [614] were the potential molecular targets of the drugs for treating diabetes mellitus. RUNX1 [395], CDKN2A [403], HSPA2 [423], hsa-mir-132-3p [615], hsa-mir-155-5p [610] and USF2 [616] have been reported to be expressed in Alzheimer’s disease. RUNX1 [449], CAV1 [472], STAT6 [468], CDKN2A [226], HSPA2 [485], hsa-mir-10b-5p [617] and HOXA5 [618] were related to the obesity. RUNX1 [511], CAV1 [534], CACNA1A [543], STAT6 [528], CDKN2A [105], hsa-mir-4516 [619], SRY (sex determining region Y) [620], FOXC1 [621] and PRDM1 [622] might offer useful information for treating hypertension. CAV1 [303], CACNA1A [307], IRS4 [317] and hsa-mir-155-5p [623] expression have been found to be altered in patients with schizophrenia. CACNA1A [348], CDKN2A [343] and hsa-mir-132-3p [624] were identified as a potential targets of neurodegenerative diseases. CACNA1A [591] can be used as a potential biomarker for an epilepsy. Increasing evidence demonstrated that CAV1 [370], hsa-mir-132- 3p [615], hsa-mir-4516 [625] and SRY (sex determining region Y) [626] have a function in Parkinson’s disease. hsa-mir-10b-5p [627] is involved in Huntington disease. hsa-mir-4516 [628] is an emerging amyotrophic lateral sclerosis biomarker. In this investigation, MYC (MYC proto-oncogene, bHLH transcription factor), MAP1LC3A, TFAP2A, GDA (guanine deaminase), MAPK13, KHDRBS2, hsa-mir-3157-5p, hsa-mir-4530, hsa-mir-4796-3p, hsa-mir-4713-5p, RELA (RELA proto-oncogene, NF-kB subunit) and REL (REL proto-oncogene, NF-kB subunit) were identified as novel therapeutic targets that might be potential biomarkers for schizophrenia. This suggests that they all play a key role in the progression of schizophrenia.

In conclusion, in the present investigation, we conducted a thorough bioinformatics and NGS data analysis of DEGs by GSE106589 data screening and identified several genes implicated in the development and progression of schizophrenia. A total of 955 genes were identified, of which MYC, FN1, CDKN2A, EEF1G, CAV1, ONECUT1, SYK, MAPK13, TFAP2A and BTK are probable core genes of schizophrenia.. This investigation reveals a series of valuable genes for further research into the non-invasive diagnosis and targeted therapy of schizophrenia.. However, bioinformatics and NGS data analyses merely indicate a general direction for further investigation. To confirm the functions of DEGs in schizophrenia, molecular biology experiments are required.

## Acknowledgement

I Gabriel E Hofman, Icahn School of Medicine at Mount Sinai, Department of Genetics and Genomics Sciences, 1 Gustave L Levy Plaza, New York, USA, very much, the author who deposited their NGS dataset GSE106589, into the public GEO database.

## Ethical approval

This article does not contain any studies with human participants or animals performed by any of the authors.

## Informed consent

No informed consent because this study does not contain human or animals participants.

## Availability of data and materials

The datasets supporting the conclusions of this article are available in the GEO (Gene Expression Omnibus) (https://www.ncbi.nlm.nih.gov/geo/) repository. [(GSE106589) https://www.ncbi.nlm.nih.gov/geo/query/acc.cgi?acc=GSE106589]

## Consent for publication

Not applicable.

## Author Contributions

B. V. - Writing original draft, and review and editing

C. V. - Software and investigation

